# Exploiting uniqueness: seed-chain-extend alignment on elastic founder graphs

**DOI:** 10.1101/2024.11.24.625039

**Authors:** Nicola Rizzo, Manuel Cáceres, Veli Mäkinen

## Abstract

Sequence-to-graph alignment is a central challenge of computational pangenomics. To overcome the theoretical hardness of the problem, state-of-the-art tools use *seed-and-extend* or *seed-chain-extend* heuristics to alignment, therefore reducing the computational resources required for the task. However, two main problems still remain: on the one hand, the daunting amount of sequencing data requires us to trade alignment accuracy with computational resources; on the other hand, current graph representations of pangenomes introduce an excessive amount of spurious recombinations.

In this paper, we implement a complete seed-chain-extend alignment workflow based on *indexable elastic founder graphs* (iEFGs), a class of graphs built from aligned sequences and supporting fast pattern matching while reducing the number of artificial recombinations. We show how to construct iEFGs from the variations to a linear reference, find high-quality seeds, and extend them using `GraphAligner`, at the scale of a telomere-to-telomere assembled human chromosome.

The main ingredient of our workflow is the use and the efficient computation of *semi-repeat-free seeds* (srf), a novel class of iEFG-based seeds introduced in this work. The amount of srf seeds is two orders of magnitude less than that of minimizers at the human chromosome level while maintaining comparable speed. Thanks to the uniqueness properties of iEFGs, we show that srf-based seeds suffice to maintain high accuracy while leveraging the speed of our tool. To further stress our point, we also implement chaining of seeds on the elastic degenerate string relaxation of the iEFG and show that only chained seeds suffice to achieve high accuracy alignments.

Our sequence-to-graph alignment tool and the scripts to replicate our experiments are available in https://github.com/algbio/SRFAligner.

## 1 Introduction

The aim of computational pangenomics is to incorporate observed genetic variants into the analysis of sequencing data [6]. This is to overcome reference bias of a unique consensus genome [12]. To tackle the challenges involved, there exist two semantically different approaches under inspection: string collection-based and graph-based pangenomes. The former builds on compressed representations of a set of haplotype sequences [22, 10, 5, 30] and the latter explicitly compacts the shared sequence fragments into graph nodes, so that paths through the graph encode the haplotypes [11, 17, 7, 31, 1, 2]. When compared to the use of a single reference sequence, both approaches need further considerations to be useful in downstream analyses: In string collections, the shared content between haplotypes is compacted without alignment information, so e.g. a sub-string match between a sequencing read and the set of haplotypes can have many occurrence locations, but only a few of them represent genetically different loci. In pangenome graphs, haplotype information can be seen as a complex tailored compression scheme for the underlying set of genomes; achieving the same compression and fast queries as in the case of a set of strings is challenging. This paper studies the graph-based approach to computational pangenomics and aims to tackle the challenge of supporting fast queries. Namely, just finding exact matches between a sequencing read and a pangenome graph is known to be as hard as finding approximate matches under the Orthogonal Vectors Hypothesis (OVH) [8, 16]. This hardness prevails even when the pangenome is represented by limited-topology objects such as elastic degenerate strings (ED-Ses) [13] and elastic founder graphs (EFGs) [9], where the former representation is a sequence of sets of strings, representing variants that can be freely recombined, and the latter is a generalization with edges specifying which strings from one set are connected to which strings in the next set, thus further restricting the possible recombinations between variants. The difficulty of overcoming the theoretical barrier in *sequence-to-graph alignment* has resulted in the introduction of heuristics like *seed-and-extend* [25, 20] and *seed-chain-extend* [21, 3], adapting their classical counterparts from the approximate string matching literature. These general heuristic frameworks are open to many optimizations, including the selection of the underlying graph representation, seeding mechanism, chaining or clustering variant, and extension algorithm.

There exists theoretical evidence that by using Maximal Exact Matches (MEMs) as seeds it is possible to obtain optimal alignments through co-linear chaining [27, 28, 15, 4]. Unfortunately, the running time of current MEM finding implementations makes that approach unpractical even for EFGs. Therefore, we introduce a tailored seeding mechanism specific to *indexable* (a.k.a. semi-repeat-free) EFGs (iEFGs) [9]. Indexable EFGs admits linear-time pattern matching, which we leverage to obtain our *semi-repeat-free seeds* (srf). Our srf seeds are not only fast to compute but also have a unique locus in the graph, thus capturing a high amount of genomic information in just a few high-quality seeds.

Furthermore, we extend `ChainX`, a practical sequence-to-sequence chaining algorithm [15], to work on the EDS relaxation of EFGs. To make a complete sequence-to-graph workflow, we feed the outcome to `GraphAligner` [25] for the final step of extending the chain into a full alignment. The overall workflow is depicted later in Figure 2.

**Figure 1.**
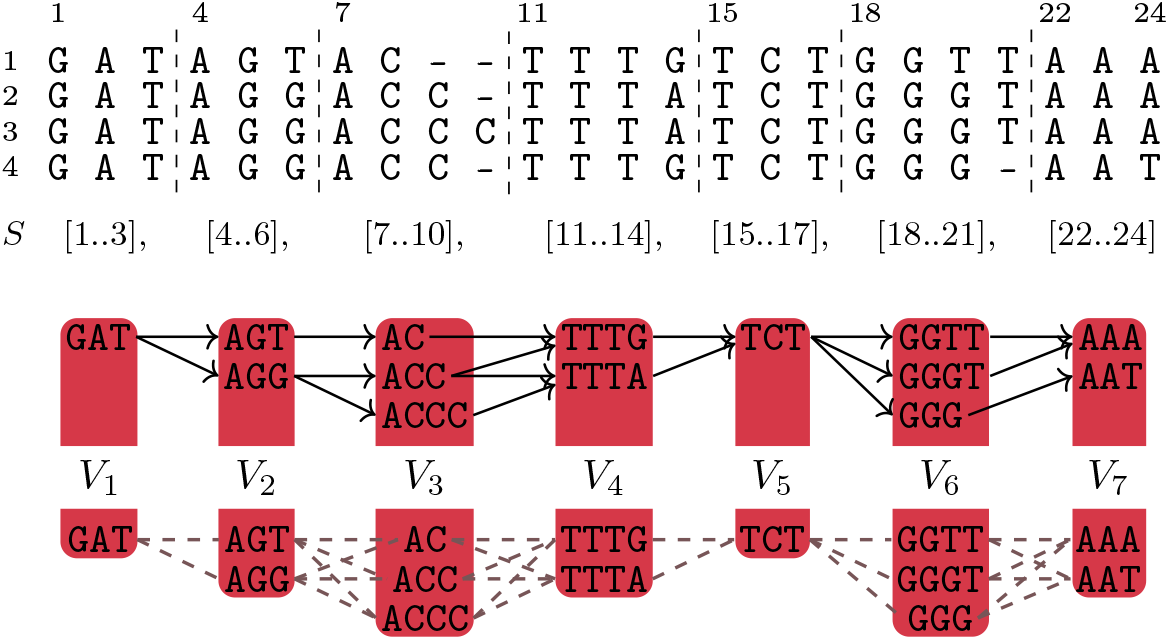
An example of an MSA[1..4, 1..24] segmented into an iEFG *G* = (*V, E*, 𝓁) of height *H*(*G*) equal to 3 and maximum node length *L*(*G*) equal to 4. Below, the relaxation of the iEFG to an elastic degenerate string *G* = (*V*, 𝓁) allowing all possible edges between consecutive blocks.

**Figure 2.**
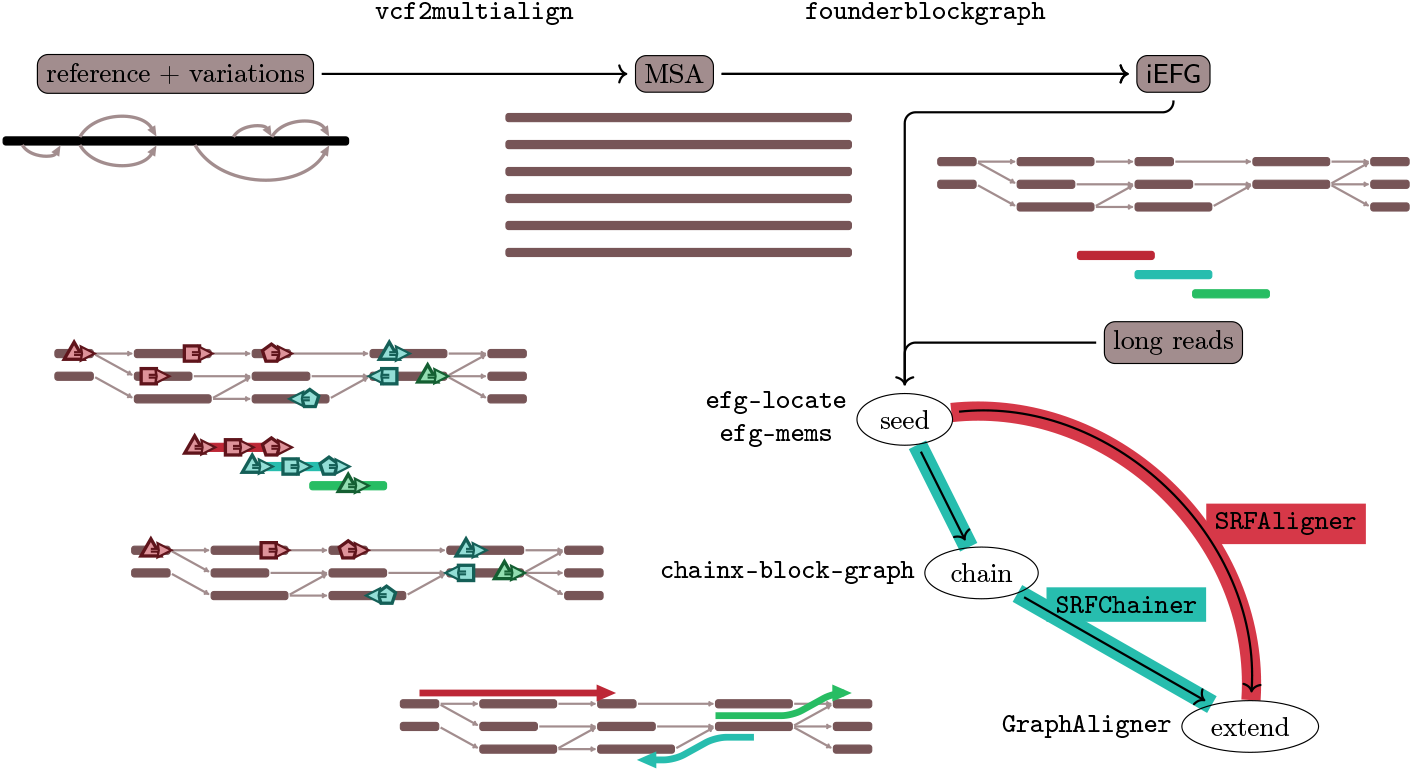
Workflow for the construction of iEFGs and the alignment of long reads: the variations (in VCF format) implicitly represent a graph that `vcf2multialign` [23] converts to a multiple sequence alignment (FASTA), that is then segmented into an iEFG (GFA). Then, the alignment is performed by finding seeds between the reads (or their reverse complement) and the graph, optionally finding the optimal chains per strand, and passing the seeds to `GraphAligner`.

### 1.1 Structure of the paper and summary of our results

In the next section, we introduce the basic concepts around iEFGs. In Section 3 we give an overview of our sequence-to-graph alignment workflow and evaluation metrics used in our experiments throughout the rest of the paper. Section 4 is devoted to the construction and validation of iEFGs at the human chromosome scale. The outcome is experimentally compared to other pangenome graph construction approaches on the human T2T-CHM13 reference and 1000 Genomes Project phased variations [24, 26, 18]. Section 5 introduces srf-based seeds and a greedy seeding algorithm to find them, and evaluates their mapping accuracy against MEMs and `GraphAligner` default minimizer seeds. While there is a small drop in alignment accuracy, the approach yields a 5x speed-up against `GraphAligner`, at the human chromosome level. Section 6 presents our `ChainX`-based [15] algorithm to chain seeds on the EDS relaxation of iEFGs. The experimental evaluation shows that this additional step reduces the number of seeds but does not significantly affect accuracy. In Section 7 we introduce a hybrid scheme that lets `GraphAligner` realign reads our approach fails to align. We compare our approaches to state-of-the-art aligners and show that our hybrid scheme achieves the highest accuracy while being 4.5 times faster than its closest competitor `GraphAligner`.

Our final product is a toolset consisting of `SRFAligner` and `SRFChainer` that implement the two alternative pipelines of the workflow depicted in Figure 2. We believe that each one of the different components of our modular workflow is of independent interest and can be repurposed independently. For this purpose, our Supplement contains thorough experiments and full details on each component, while this main paper focuses on the overall workflow.

## 2 Preliminaries

Given positive integers *x* ≤ *y*, we denote range {*x, x* + 1, …, *y*} as [*x*..*y*]. Given a finite alphabet Σ, we say that *Q* = *Q*[1]*Q*[2] · · · *Q*[*n*] ∈ Σ^*n*^ is a string of length *n* over Σ, and also that |*Q*| = *n*. Then, we define Σ^∗^ as the set of all strings over Σ and Σ^+^ = Σ *\* {*ε*}, where *ε* is the string of length 0. We also define *Q*[*i*..*j*] as the concatenation *Q*[*i*]*Q*[*i* + 1] · · · *Q*[*j*] of the symbols of *Q* from the *i*-th to the *j*-th. String *P* is then a *substring* of *Q* if *P* = *Q*[*i*..*j*] for some *i, j* ∈ [1..|*Q*|]: if *P≠ Q* then we say that *P* is *proper* ; if *i* = 1 then *P* is a *prefix* of *Q* and we also write *Q*[..*j*]; if *j* = |*Q*| it is a *suffix* and we also write *Q*[*i*..]. Further, we say that *i* is an *occurrence* of *P* in *Q*, and if *P* [*x*..*y*] = *Q*[*a*..*b*] then ([*x*..*y*], [*a*..*b*]) is an *(exact match) anchor* between *P* and *Q*.

In this paper, we deal with graphs built from the segmentation of multiple sequence alignments, represented as a matrix MSA[1..*m*, 1..*n*] ∈ (Σ ∪ {−})^*m*×*n*^ representing *m* sequences (the rows MSA[*i*, 1..*n*] for *i* ∈ [1..*m*]) aligned into *n* positions (the columns). Alphabet Σ is expanded with the special *gap character* − ∉ Σ representing insertions and deletions: given an aligned sequence MSA[*i*, 1..*n*] we can retrieve the original sequence spell(MSA[*i*, 1..*n*]) by removing the gaps. We define a *segmentation S* of MSA[1..*m*, 1..*n*] as a partition of columns [1..*n*] in *b* segments [1 = *x*_1_..*y*_1_], …, [*x*_*b*_..*y*_*b*_ = *n*] such that *y*_*i*_ = *x*_*i*+1_ − 1 for *i* ∈ [1..*b* − 1]. We consider the following graph objects, see also Figure 1.

**Definition 1** (elastic founder graph [9]). *Let* MSA[1..*m*, 1..*n*] ∈ (Σ∪{−})^*m*×*n*^ *be a multiple sequence alignment. Given segmentation S* = [*x*_1_..*y*_1_], …, [*x*_*b*_..*y*_*b*_], *the* elastic founder graph *(EFG)* induced by *S is a vertex-labeled graph G* = (*V, E*, 𝓁), *where* 𝓁 : *V* → Σ^+^ *assigns a non-empty* label *to every node, and the set of nodes V is partitioned into blocks V*_1_, …, *V*_*b*_ *such that* |{ 𝓁(*v*) : *v* ∈ *V*_*k*_ }| = |*V*_*k*_| *(unique strings in a block) and*

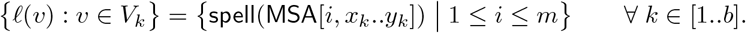

*Moreover, it holds that* (*v, w*) ∈ *E if and only if there exists k* ∈ [1..*b* − 1] *and i* ∈ [1..*m*] *such that v* ∈ *V*_*k*_, *w* ∈ *V*_*k*+1_, *and* spell(MSA[*i, x*_*k*_..*y*_*k*+1_]) = 𝓁(*v*)𝓁(*w*).

If 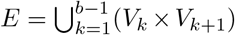, that is all nodes in *V*_*k*_ are connected to all nodes in *V*_*k*+1_ for each *b* ∈ [1..*b*], then the EFG is an elastic degenerate string (EDS) [14], and we can omit edge set *E*. Moreover, given an EFG, we define its EDS relaxation as the corresponding EDS over *V*_1_, …, *V*_*b*_.

Given an elastic founder graph *G* = (*V, E*, 𝓁), for each node *v* ∈ *V* we denote as ∥*v*∥ the node label length |𝓁(*v*)|. We define height 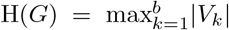 as the maximum number of nodes in a block, and L(*G*) = max_*v*∈*V*_ ∥*v*∥ as its maximum node label length. We can extend 𝓁 to label paths in the graph: given a *u*_1_*u*_*k*_-path *P* = *u*_1_ · · · *u*_*k*_, with (*u*_*i*_, *u*_*i*+1_) ∈ *E* for *i* ∈ [1..*k* − 1], we define 𝓁(*P*) = 𝓁(*u*_1_) · · · 𝓁(*u*_*k*_); for an edge (*u, v*) ∈ *E*, we call 𝓁(*uv*) its *edge label*. Then, we say that some string *Q* ∈ Σ^+^ *occurs* in *G* if there exists path *P* in *G* such that *Q* is a substring of 𝓁(*P*). More specifically, we indicate with triple (*i, u*_1_ · · · *u*_*k*_, *j*) the *subpath* of *P spelling Q* starting from position *i* ∈ [1..∥*u*_1_∥] in *u*_1_ and ending in position *j* ∈ [1..∥*u*_*k*_∥] in *u*_*k*_, that is, *Q* = 𝓁(*u*_1_)[*i*..]𝓁(*u*_2_ · · · *u*_*k*−1_)𝓁(*u*_*k*_)[..*j*]. The *length* of subpath (*i, u*_1_ · · · *u*_*k*_, *j*) is then |*Q*|. We say that *Q* occurs *starting from the beginning of u*_1_ if *i* = 1; *Q* occurs *starting inside u*_1_ if *i* ∈ [2..∥*u*_1_∥]. Moreover, if substring *Q*[*x*..*y*] is spelled by subpath (*i, P* = *u*_1_ · · · *u*_*k*_, *j*), we say that pair ([*x*..*y*], (*i, P, j*)) is an *(exact match) anchor* between *Q* and *G*, and if *k* = 1 we say that it is a *node anchor*.

Unfortunately, EFGs are still hard to index for subquadratic-time pattern matching [9], and an additional property such as the following one is required.

**Definition 2** (semi-repeat-free indexability property [9]). *An EFG G* = (*V, E*, 𝓁) *is* semi-repeat-free *or* indexable *(denoted as iEFG) if each subpath* (*i, u*_1_…*u*_*k*_, *j*) *spelling* 𝓁(*v*) *for v* ∈ *V*_*i*_ *is such that i* = 1 *and u*_1_ ∈ *V*_*i*_, *that is*, 𝓁(*v*) *only occurs from the beggining of nodes in block V*_*i*_.

## 3 Workflow overview and aligner evaluation

In this paper, we implement the different steps of the workflow illustrated in Figure 2 to align long reads to an iEFG.

We measure the *accuracy* of the aligners as the numbers of correctly aligned reads divided by the total number of input reads. Sequence-to-graph aligners align each read to a subpath of the graph: we call such a path the *reported subpath* and the string spelled by this the *reported sequence*. If there are multiple reported alignments per read, we take the primary one (the first) and discard the secondary alignments (the rest) for `GraphAligner`-based tools, and for other tools we take the alignment with the longest reported subpath^1^. We define the correctness of an alignment with three different parameterized criteria introduced by Ma et al. [21]:

### Path accuracy

Given a parameter 0 *< δ* ≤ 1 we say that an alignment is correct if the base-level overlap between the reported subpath and the true subpath is at least *δ* times the true subpath length. This is a generalization of the widely used 10% overlap criteria, that is, *δ* = 0.1 [19, 25, 4].

### Truth ED accuracy

Given 0 *< σ*_truth_ ≤ 1 we say that the alignment is correct if the Levenshtein distance between the reported sequence and true sequence is at most *σ*_truth_ times the length of the true sequence.

### Read ED accuracy

Given 0 *< σ*_read_ ≤ 1 we say that the alignment is correct if the Levenshtein distance between the read and the reported sequence is at most *σ*_read_ times the read length.

The ground truth is generated by choosing a random path through the graph in question and sampling long reads from it using `BadRead` simulator [32]. The simulator, unless stated otherwise, is run with an error level of 5% under its Oxford Nanopore R10.4.1 model, target coverage of 30x, and target length of 15000bp.

The machine used in the experiments runs on an Intel(R) Xeon(R) CPU E7-4830 v3 @ 2.10GHz and 1.5 TB of RAM. SSD storage space is used in all experiments except in the graph construction experiments, where a large amount of disk space was needed and HDD storage space was used. Unless stated otherwise, each tool is executed using 64 threads, and the aligners do not use additional disk space nor reuse indexes for the graph or the reads.

## 4 Elastic founder graph construction

We built chromosome iEFGs from the recent T2T-CHM13 reference genome and the variation from the 1000 Genomes Project (1KGP) recalled and phased on this reference [24, 26, 18]. For our experiments, we use the chromosome 22 iEFG. The various engineering choices required to make this feasible on top of the earlier theoretical construction algorithms are detailed in the Supplement; for an overview, see Figure 3. The Supplement also contains a thorough study of the uniqueness of a human pangenome, as well as a comparison to the `vg` toolkit [12], summarized in Table 1. We tested `vg` graphs built from a VCF file and from the corresponding MSA obtained with `vcf2multialign` [23]. In both cases, graphs produced by `vg` contain several orders of magnitude more paths than the iEFG, showing that iEFG effectively reduces the number of spurious recombinations. Furthermore, the accuracy of the alignment is not significantly impacted on the iEFG (see Supplement). However, we observe that the number of minimizer seeds found in the iEFG is 5-6 times more, increasing the computational resources used (see Supplement) and motivating our further study of iEFG-specific seeds.

**Table 1:**
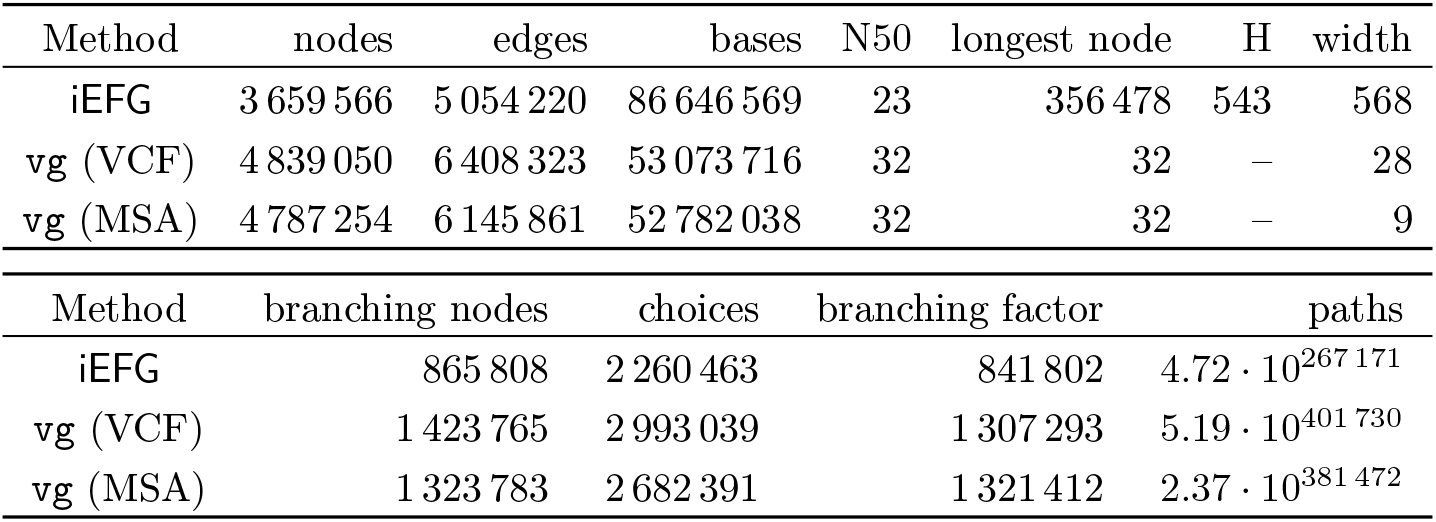
Features of the chromosome 22 graphs generated from the T2T-CHM13 reference and the 1KGP phased variations. Height H is defined as the largest number of nodes in an iEFG block; width is the size of the smallest set of paths covering the graph nodes; choices is the cumulative degree of branching nodes; branching factor is the largest number of branching nodes encountered in any path; paths is the total number of maximal paths expressed in the graph, represented in scientific notation rounded to the 2nd decimal.

**Figure 3.**
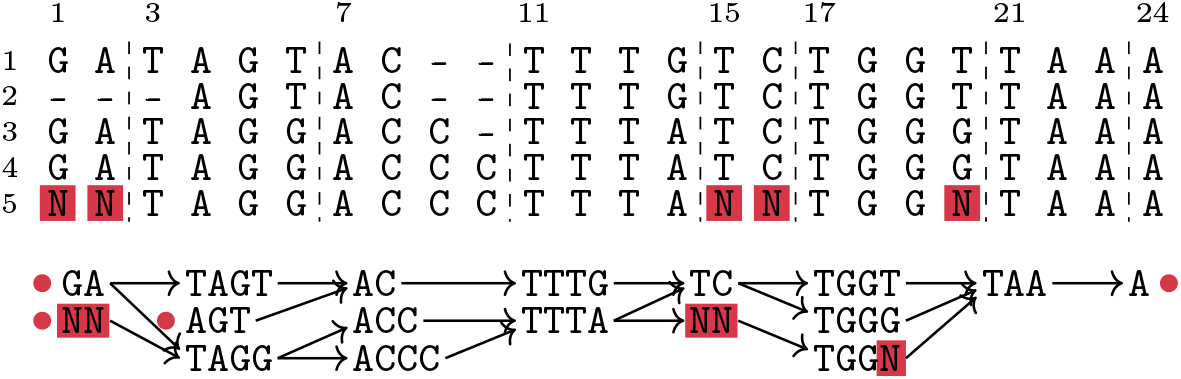
Some of the engineering techniques applied to the practical construction of iEFGs. For example, we can assume each row starts and ends with a unique character (marked with a dot), allow empty segments at the beginning and end of sequences, and ignore `N`s by considering them as unique. Pattern matching results and additional properties (see Supplement) still hold if we do not query patterns containing `N`s.

## 5 Exact pattern matching and unique seeds

Indexable EFGs admit linear-time exact pattern matching by indexing (a variation of) the following string spelled by the concatenation of edge labels [29]:

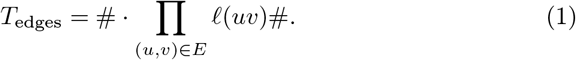

During pattern matching, if a pattern spans more than two nodes, special attention must be given to the last few blocks matching the string: in [29, Sections 3 and 6], this is solved by complex algorithms requiring several advanced data structures. In the Supplement, we give a simplified solution that has a slower worst-case running time but is amenable to a practical implementation that extends to a new seeding technique. This yields the following result.

### Theorem 1

(Theorem 4 in the Supplement). *Let G* = (*V, E*, 𝓁) *be an iEFG. There is an algorithm that, after a preprocessing of G in O* |*T*_*edges*_| *time (Equation* (1)*), answers whether Q* ∈ Σ^+^ *occurs in G in O*(|*Q*| + min(|*Q*|, L(*G*))^2^ + H(*G*)^2^) *time, where* L(*G*) *is the longest node length and* H(*G*) *is the maximum block size. In the positive case, the algorithm also reports a match of Q in G*.

In order to exploit the uniqueness properties of iEFGs as a seeding technique, we formalize the following type of seed.

**Definition 3** (Semi-repeat-free seed). *An anchor* ([*x*..*y*], (*i, P* = *u*_1_ · · · *u*_*k*_, *j*)) *between iEFG G* = (*V, E*, 𝓁) *and pattern Q* ∈ Σ^+^ *is a* semi-repeat-free seed *if subpath* (*i, P, j*) *spans at least a full node, defined as follows: k >* 2, *or if k* = 1 *then i* = 1 *and j* = ∥*u*_*k*_∥, *or k* = 2 *and at least one of conditions i* = 1 *and j* = ∥*u*_*k*_∥ *hold*.

We show a greedy approach to finding a subset of semi-repeat-free seeds. In our algorithm, we apply exact pattern matching from Theorem 1 as follows:

1. We use Theorem 1 to find a long occurrence of a suffix *Q*[*y*..] in *G*;
2. If the exact match (*Q*[*y*..], (*i, P, j*)) found in the previous step spans at least a full node, we report it as a semi-repeat-free seed, otherwise we report no match and continue the search on *Q*[1..*y* − 1], stopping when *y* − 1 = 0.

We now explain why we expect this subset of semi-repeat-free seeds to be of high quality in practice. Consider any exact match anchor ([*x*..*y*], (*i, P* = *u*_1_ · · · *u*_*k*_, *j*)) between *Q* and *G* with *P* spanning four or more full nodes, that is, *k* ≥ 4, *i* = 1, and *j* = ∥*u*_*k*_∥. The previous greedy approach will report a semi-repeat-free seed involving *Q*[*x*..*x* + |𝓁(*u*_1_ … *u*_*k*−2_)|] and *u*_1_ due to the properties of iEFGs.

We implemented the above solutions to pattern matching and seeding in our tool `efg-locate`. We experimentally tested alignment accuracy over the default minimizers of `GraphAligner` and Maximal Exact Matches (MEMs) as seeds. While MEMs are too slow for practical use, srf-based seeds outperform minimizer seeds and retain most of the accuracy, as seen in Table 2 (also in the Supplement).

**Table 2:**
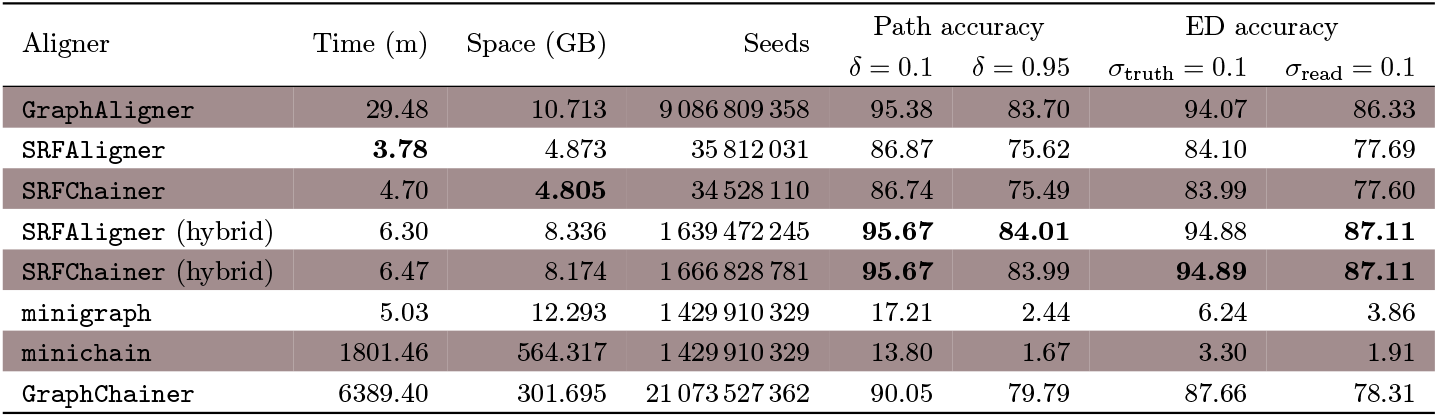
Comparison of aligner performance and accuracy of simulated reads for the chromosome 22 iEFG built from the T2T-CHM13 + 1KGP dataset.

| Aligner | Time (m) | Space (GB) | Seeds | Path accuracy |  | ED accuracy |  |
| --- | --- | --- | --- | --- | --- | --- | --- |
| | | | | $\delta = 0.1$ | $\delta = 0.95$ | $\sigma_{\text{truth}} = 0.1$ | $\sigma_{\text{read}} = 0.1$ |
| **GraphAligner** | 29.48 | 10.713 | 9 086 809 358 | 95.38 | 83.70 | 94.07 | 86.33 |
| **SRFAligner** | **3.78** | 4.873 | 35 812 031 | 86.87 | 75.62 | 84.10 | 77.69 |
| **SRFChainer** | 4.70 | **4.805** | 34 528 110 | 86.74 | 75.49 | 83.99 | 77.60 |
| SRFAligner (hybrid) | 6.30 | 8.336 | 1 639 472 245 | **95.67** | **84.01** | 94.88 | **87.11** |
| **SRFChainer (hybrid)** | 6.47 | 8.174 | 1 666 828 781 | **95.67** | 83.99 | **94.89** | **87.11** |
| minigraph | 5.03 | 12.293 | 1 429 910 329 | 17.21 | 2.44 | 6.24 | 3.86 |
| minichain | 1801.46 | 564.317 | 1 429 910 329 | 13.80 | 1.67 | 3.30 | 1.91 |
| GraphChainer | 6389.40 | 301.695 | 21 073 527 362 | 90.05 | 79.79 | 87.66 | 78.31 |

## 6 Chaining on elastic founder graphs

We show how to chain a set of exact match node anchors on an iEFG by adapting the sequence-to-sequence co-linear chaining formulation introduced by Jain et al. [15]. To obtain an efficient chaining solution, we break down the srf-based seeds into node anchors, so that co-linear chaining captures the desired alignment metrics [27, 15], and we relax the topology of the iEFG into its EDS relaxation (see Figure 1).

Given a set *A* of node anchors, chaining consists of selecting an ordered subset of *A* (a *chain*) that is compatible with an alignment. There exist many different versions of co-linear chaining in the literature. What defines a chaining problem are the following two concepts:

- *Precedence between anchors*: This defines when an anchor can be *chained* to the next anchor in a chain. In our case anchor *A*_*p*_ = ([*x*_*p*_..*y*_*p*_], (*i*_*p*_, *u*_*p*_, *j*_*p*_)) *precedes* anchor *A*_*q*_ = ([*x*_*q*_..*y*_*q*_], (*i*_*q*_, *u*_*q*_, *j*_*q*_)) if *x*_*p*_ *< x*_*q*_, *y*_*p*_ *< y*_*q*_ (precedence in the query), and there is a *u*_*p*_*u*_*q*_-path in *G* or *u*_*p*_ = *u*_*q*_ and *i*_*p*_ *< i*_*q*_ and *j*_*p*_ *< j*_*q*_ (precedence in the graph). The previous definition of graph precedence works for general graphs and it has been used by state-of-the-art tools [21, 4]. In the case of EDSes, we can further simplify this definition to a block-based computation. Namely, there exists a *u*_*p*_*u*_*q*_-path if and only if the block of *u*_*p*_ precedes the block of *u*_*q*_. See the Supplement for a detailed description and Figure 4 for an example.
- *Chaining cost* : This defines the objective function to optimize to find an optimal chaining. We use the same objective function as in [15], which provides a theoretical connection to the Levenshtein distance in the linear case. We aim to minimize the *cost* of a chain *A*_1_, …, *A*_*c*_ defined as

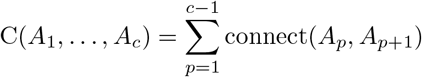

where connect(*A*_*p*_, *A*_*p*+1_) = g(*A*_*p*_, *A*_*p*+1_) + o(*A*_*p*_, *A*_*p*+1_) with

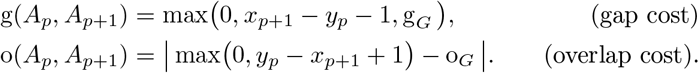

The graph gap g_*G*_ is the shortest length of a (*i*_*p*_, *u*_*p*_, *j*_*p*_)-(*i*_*q*_, *u*_*q*_, *j*_*q*_) subpath in *G*^2^ minus the lengths of the two anchors, and the graph overlap o_*G*_ is 0 if *u*_*p*_*≠ u*_*q*_ and max(0, *j*_*p*_ − *i*_*p*+1_ + 1) otherwise. Additionally, we add one initial anchor that precedes all other anchors and one final anchor that is preceded by all other anchors. The connect function on these two anchors defines global and semi-global chaining modes. See the Supplement for details and Figure 4 for an example.

**Figure 4.**
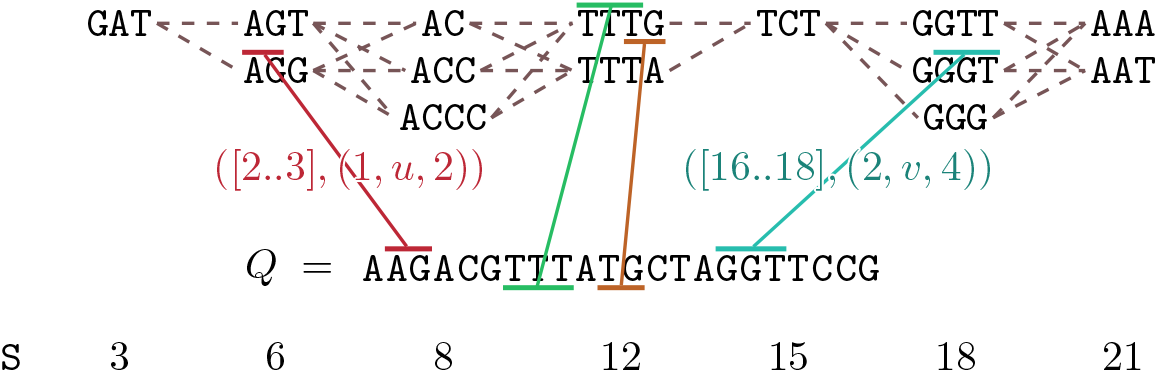
An EDS *G* = (*V*, 𝓁) partitioned into blocks *V*_1_, …, *V*_*b*_, a pattern *Q*, and a chain of co-linear node anchors between *Q* and *G*. The cost of connecting consecutive anchors in the chain, defined as the maximum gap plus the absolute overlap difference, can be computed in constant time by preprocessing *G* for the constant-time computation of node block indices and distance between two EDS nodes.

Co-linear chaining can be solved by a simple dynamic programming approach with the following recurrence. Let *A*_1_, …, *A*_*n*_ be the anchors in *A* sorted by *x*_*p*_ (starting position in *Q*). Then, the cost of an optimal chain ending with *A*_*q*_ is

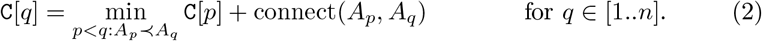

This dynamic programming formulation can be solved in *O*(*n*^2^) time provided that we can check precedence and compute the connect function in constant time as follows. Indeed, we can preprocess EDS *G* = (*V*, 𝓁) partitioned into blocks *V*_1_, …, *V*_*b*_ in linear time to answer the following queries:

- *Precedence in constant time*: It suffices to precompute table B such that B[*u*] = *i* is the block of *u* ∈ *V*_*i*_.
- *Connect in constant time*: It suffices to additionally precompute table S such that 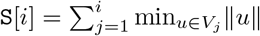 is the length of the shortest path from the start of block *V*_1_ to the end of block *V*_*i*_, since we consider the EDS relaxation of *G* (see Figure 4). Then, the length of the shortest (*i*_*p*_, *u*_*p*_, *j*_*p*_)-(*i*_*q*_, *u*_*q*_, *j*_*q*_) subpath in *G* is (∥*u*_*p*_∥ − *i*_*p*_ + 1) + S[B[*u*_*q*_] − 1] − S[B[*u*_*p*_]] + *j*_*q*_ if *u*_*p*_*≠ u*_*q*_, otherwise it is *j*_*q*_ − *i*_*p*_ + 1.

Our final ingredient is the following inspection of preceding anchors similar to one implemented in `ChainX` [15]. In Equation (2), we restrict the search of anchors with gap distance *x*_*q*_ − *y*_*p*_ − 1 ≤ *B* (with *B* an initial guess) in *Q* and previous anchors are not inspected. If C[*n*] *> B*, we update *B* to *B* · *α* (a ramp-up factor) and recompute values C[1..*n*] until C[*n*] ≤ *B*. In the Supplement, we prove that under restricted conditions on the anchors we obtain the following result.

### Theorem 2

(Theorem 8 in the Supplement). *Let Q* ∈ Σ^+^ *be a pattern and G* = (*V*, 𝓁) *an EDS preprocessed for constant-time precedence and connect queries. Given n anchors between Q and G, we can find an optimal chain of cost* OPT *in O*(*n* · OPT + *n* log *n*) *average-case time, assuming that n* ≤ |*Q*|, *the anchors do not overlap in the query, and the anchor endpoints are uniformly distributed*.

Semi-repeat-free anchors obtained with the greedy method of Section 5 respect the conditions of Theorem 2. We implemented co-linear chaining on the EDS relaxation of an iEFG in tool `chainx-block-graph`. Our experiments show that performing chaining on srf-based seeds maintains comparable speed while reducing the number of final seeds and peak memory consumption required by `GraphAligner` (see Table 2 and Supplement).

## 7 Final comparison and discussion

Our final toolset consists of `SRFAligner` and `SRFChainer` which implement the two alternative full pipelines of the workflow depicted in Figure 2, the former without chaining and the latter with chaining. In addition, they support a *hybrid* mode, where poorly aligned reads^3^ are re-aligned with the default setting (minimizer seeds) of `GraphAligner`. Table 2 shows the comparison of different modes of `SRFAligner` and `SRFChainer` to default `GraphAligner` [25] as well as to `minigraph` [20], `minichain` [3], and `GraphChainer` [21]. Column *seeds* is the number of seeds provided to the extension phase.

The high resource usage of `minichain` and `GraphChainer` is explained by the high width of iEFG (see Table 1), as their performance is heavily dependent on this parameter. The low accuracy of `minigraph` and `minichain` on complex pangenome graphs with short node labels is a known limitation reported at https://github.com/lh3/minigraph#limitations.

The hybrid modes of `SRFAligner` and `SRFChainer` offer the best performance and accuracy with 4.5x speed-up against default `GraphAligner`, while retaining the vast majority of correct alignments.

We remark that srf-based seeds are not only relevant for iEFGs but promising for all pangenome representations, as they characterize the uniqueness of genomic sequences. We plan to study how to project our seeds to other representations.

## Footnotes

1 We observed better accuracy with this method for other tools, as reported in [21].

2 This subpath is of the form (i_p_, P = u_p_· · · u_q_, j_q_) such that P is a u_p_u_q_-path in G.

3 Alignments with `GraphAligner` identity of ≤ 90% or read coverage of < 50%.

