## Supplement for "Exploiting uniqueness: seed-chain-extend alignment on elastic founder graphs"

**A Characterization of iEFGs** **18**

**B Exact pattern matching in iEFGs** **19**

**C Full results of experimental tests** **26**

**D Commands used in the experiments** **28**

#### 2 Preliminaries

See the main paper for basic definitions.

To index iEFGs and EDSes, we use the well-known machinery of text indexing and bitvectors behind the FM-index [6]. For a bitarray $B[1...n] \in \{0,1\}^n$, we use the well-known *rank* and *select* queries over $B$: given a position $i \in [1..n]$ rank answers the number of ones in $B[1..i]$, whereas given a rank position $j \in [1..\text{rank}(n, B)]$, selects return the position of the $j$-th one in $B$. The bit vectors used can be preprocessed to answer these queries in constant time [11]. Besides, we preprocess a text $T$ for *backward pattern matching* based on the suffix-array: we consider the suffixes $T[i..n+1]$ of $T\$$ in their lexicographical order, with $\$ \notin \Sigma$ a special terminator character; we can search for the occurrences of pattern $Q \in \Sigma^+$ in $T$ by iteratively considering the range $[\ell..r]$ in the list of sorted suffixes corresponding to suffix $Q[i..]$; importantly, this range corresponds to all occurrences of $Q[i..]$ in $T$. The main primitive of backward search is then operation LeftExtend$(T, [\ell..r], c)$, that given $c \in \Sigma$ and interval $[\ell..r]$ corresponding to string $Q$ returns the interval corresponding to string $c \cdot Q$.

The semi-repeat-free property and the definition of elastic founder graph imply the following useful properties, as described and used in [5].

**Lemma 1** (semi-repeat-free uniqueness properties)**.** *Given an* iEFG $G = (V, E, \ell)$*:*

- *any two distinct nodes* $u \neq v$, $u, v \in V$*, cannot have the same node label, i.e.* $\ell(u) \neq \ell(v)$*;*
- *no* $\ell(u)$ *with* $u \in V$ *can be a proper suffix of* $\ell(v)$ *for some* $v \in V$*;*
- *all occurrences of* $\ell(uv)$ *in* $G$ *with* $u, v \in V$ *start from the beginning of* $u$*.*

Regarding the construction of iEFGs, Equi et al. in [5] proved that all segments resulting in an iEFG are fully described by values $f(x)$: given $x \in [1..n]$, $[x..f(x)]$ is the smallest segment starting from $x$ and such that the strings in the segment occur in the MSA only starting from column $x$; $f(x) = \infty$ if such segment does not exist; then, any segment $[x..y]$ is valid if and only if $y \geq f(x)$ (see Figure 1). Using values $f()$, an $O(mn)$-time solution to find the optimal segmentation minimizing the maximum segment length, which approximates the maximum node length $L(G)$ of the resulting iEFG, was proposed and implemented by Rizzo et al. [15] in tool `founderblockgraph`.

**Observation 1** (iEFG recombination model)**.** *Consider an* iEFG $G = (V, E, \ell)$ *generated from the segmentation of some* MSA$[1..m, 1..n]$ *and let* $P_1, \ldots, P_m$ *be the graph paths of* $G$ *corresponding to rows* MSA$[1, 1..n]$*,* $\ldots$*,* MSA$[m, 1..n]$*. For any path* $P = uvw$ *of length* 3 *we have that if* $P$ *is not a substring of some* $P_i$*,* $i \in [1..m]$*, then it spells string* $\ell(uvw)$ *where* $\ell(uv)$*,* $\ell(vw)$ *occur in some of the original sequences and* $\ell(v)$ *is a semi-repeat-free substring of* MSA$[1..m, 1..n]$*: we call this a* semi-repeat-free recombination*. Thus, any substring spelled by the paths of* $G$ *either occurs in the original sequences or is the result of non-overlapping semi-repeat-free recombinations of the sequences.*

#### 4 Elastic founder graph construction

The current theory on the segmentation of aligned sequences into iEFGs presents a few practical challenges on real datasets:

1. long runs of gap symbols in the MSA constrain heavily the possible segmentations, since segments whose spelling is the empty string for some row $i$, that is, spell(MSA$[i, x..y]$) $= \varepsilon$, are not allowed [16, Assumption 1];

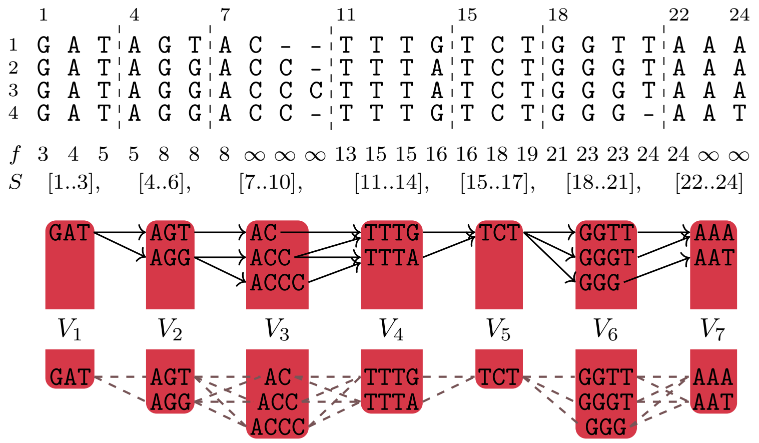

Figure 1: An example of an $\mathsf{MSA}[1..4, 1..24]$ segmented into an $\mathsf{iEFG}$ $G = (V, E, \ell)$ of height $H(G)$ equal to 3 and maximum node length $L(G)$ equal to 4. Values $f()$ fully describe the possible segments resulting in an $\mathsf{iEFG}$: $f(x)$ is equal to the minimum $y$ such that segment $[x..y]$ contains unique substrings occurring only from column $x$. Below, the relaxation of the $\mathsf{iEFG}$ to an elastic degenerate string $G = (V, \ell)$ allowing all possible edges between consecutive blocks.

2. if, for some rows $i, i' \in [1..m]$, we have that $\mathsf{spell}(\mathsf{MSA}[i, 1..n])$ is a proper suffix of some $\mathsf{spell}(\mathsf{MSA}[i', 1..n])$, then there is no valid segmentation, since for any two nodes $u, v \in V_1$ in the first $\mathsf{iEFG}$ block $\ell(u)$ cannot be a proper suffix of $\ell(v)$ (Lemma 1);

3. genomic data can contain missing or masked nucleotides—usually, letter `N` is used for this purpose [3]—and if the longest run of `N` has length $L$, then no node label can spell string $\mathtt{NN}\cdots\mathtt{N}$ of length $k < L$, further constraining the possible segmentations.

We can mitigate challenge 1 and solve challenges 2 and 3 with the following strategies:

(a) we assume implicitly that every row $\mathsf{MSA}[i, 1..n]$ starts and ends with unique characters that do not occur anywhere else, i.e. $\mathsf{MSA}[i, 1] = \$_i$ and $\mathsf{MSA}[i, n] = \#_i$; after computing the semi-repeat-free segmentation (and before obtaining the $\mathsf{iEFG}$) we remove all occurrences of $\$_i$ and $\#_i$;

(b) we allow segments producing the empty string $\varepsilon$ if they correspond to an initial or ending run of gaps for some row;

(c) we also consider each occurrence of some missing nucleotide $\mathsf{MSA}[i, j] = \mathtt{N}$ as an occurrence of a unique character $\mathtt{N}_{i,j}$ that does not occur anywhere else in the MSA.

See Figure 2 for an example.

Intuitively, strategies (a) and (b) do not affect the indexability of the resulting $\mathsf{EFG}$, since we can prepend and append the unique characters to the label of all sources and sinks of the graph, respectively: any pattern $Q \in \Sigma^+$ occurs in this modified graph if and only if it occurs in the original. Strategy (c) results in a graph that is *partially indexable*: we can still index the graph for pattern matching, provided that the queries will not contain character `N`. The recombination model (Observation 1) is also only partially affected, as consecutive graph edges of nodes that do not contain `N` characters still share a semi-repeat-free string.

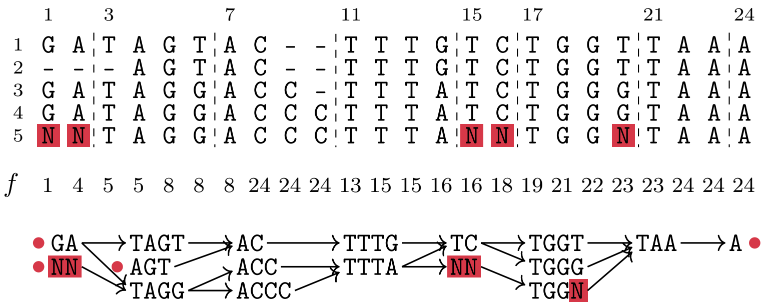

Figure 2: In the construction of iEFGs, we can assume each row starts and ends with a unique character (strategy (a)), allow empty strings in the beginning and end of the segmentation (strategy (b)), and ignore Ns by considering them as unique (strategy (c)). Sources and sinks of the iEFG are marked with a dot. Pattern matching and the recombination property (Observation 1) still hold if we do not query or consider patterns containing Ns.

We implemented the above strategies into tool `founderblockgraphs`. When a large MSA is given in input, the bottleneck of the construction is in the computation of the suffix tree of the MSA: to reduce the time and space used, we developed a heuristic construction solution that approximates the values of $f$ by considering only $M$ (a user-defined parameter) MSA rows at a time and takes the maximum between the $f(x)$ values for each column $x \in [1..n]$ to compute the segmentation; if the resulting graph is not indexable, the invalid segments are iteratively removed until the semi-repeat-free property holds, sacrificing the optimality of the segmentation. See Appendix A for the theoretical details on how we can check whether a given EFG is indexable in polynomial time.

#### 4.1 Experimental test: uniqueness of the human genome

The effectiveness of iEFGs is highly dependent on the uniqueness of the MSA in input: for example, if any of the input sequences contains long exact repetitions, no segment $[x..y]$ contained in these regions can be picked for a segmentation. To evaluate the quality of an iEFG, we can use the following contiguity metric based on the widely used N50 statistic for assembly quality [2]. As opposed to the normal use of the metric, we are interested in how much the graph is covered by considering short node labels, instead of long contigs[1].

**Definition 1** ($\lceil$N50$\rceil$ metric)**.** *For an iEFG $G = (V, E, \ell)$, we define $\lceil \mathrm{N50} \rceil(G)$ as the minimum length $L \in \mathbb{N}$ such that at least 50% of the graph bases are contained in node labels of length* at most $L$. *Analogously, given $X \in [0..100]$, we define $\lceil \mathrm{NX} \rceil(G)$ as the minimum length $L$ such that at least $(100 - X)$% of the graph bases are contained in labels of length at most $L$.*

To study the uniqueness of the human genome in terms of our indexing property, we built chromosome iEFGs from the recent T2T-CHM13 reference genome and the variation from the 1000 Genomes Project (1KGP) recalled and phased on this reference [13, 14, 10]. The reference contains complete assemblies of the human cell line CHM13 and the 1KGP collected variation from more than 2000 samples. As an optimistic baseline, we first computed the semi-repeat-free segmentation of the individual chromosomes, obtaining 24 iEFGs consisting of a linear path. The maximum segment length of the chromosome segmentation ranges from 9 483 to 1 579 331, from 0.006% to

[1]In practice, we did not see a difference between the N50 and $\lceil$N50$\rceil$ metrics, so in the main paper we just write N50.

1.832% of the full length of the sequences, showing that there are inherently challenging regions for the semi-repeat-free property. However, the ⌈N50⌉ metric is equal to 15 or 16 for all chromosomes except Y, at value 937, and the values of ⌈N5⌉ are in the range 38–8 002, showing that the majority of each sequence is segmented in unique labels of length in the order of tens or hundreds, against a sequence containing millions of bases. See Figure 3 for a visualization of the segment length distribution for the chromosome 22 iEFG segmentation. The full table of results and visualization of ⌈N50⌉ and ⌈N5⌉ values appear in Table 7 and Figure 9 in Appendix C.

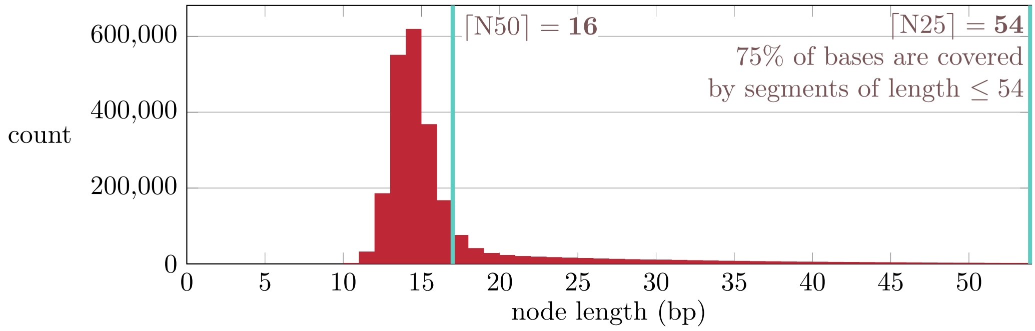

Figure 3: Histogram of the segment lengths for the semi-repeat-free segmentation of chromosome 22 (T2T-CHM13). The chromosome is partitioned in unique substrings minimizing the maximum substring length.

We built the chromosome 22 iEFG including the 1KGP variations as follows. Using tool `vcf2multialign` [12], we obtain an MSA consisting of 5009 aligned sequences representing the T2T-CHM13 reference plus the 2504 phased diploid samples. Since the input VCF file contains variations that are not fully consistent with 5008 paths of the implicit VCF graph, we performed quality control on the output of `vcf2multialign` as follows. We used `edlib` [17] to align each sequences found by `vcf2multialign` to the corresponding output given by `bcftools consensus` [4]: the average Levenshtein edit distance between the sequences is 40 and the maximum is 182, showing very high similarity. Finally, we created the corresponding iEFG with `founderblockgraphs`'s heuristic construction, setting $M = 250$. This final step took the overwhelming majority of the computational resources, approximately 6 days and 500 GB of memory. Table 1 shows the uniqueness metrics for the chromosome 22 iEFG compared to the segmentation of the same chromosome (i.e. only the reference): the comparison of ⌈NX⌉ values shows that the majority of the variation added is contained in short sequences. The chromosome 22 iEFG will be used in later sections for our experiments.

Table 1: Comparison of uniqueness statistics between the semi-repeat-free segmentation of the linear T2T-CHM13 chromosome 22 reference and the iEFG built from the same reference plus the 1KGP phased variations using `vcf2multialign` and `founderblockgraph`.

| Graph | Bases | Semi-repeat-free segmentation | | | | |
|---|---|---|---|---|---|---|
| | | ⌈N50⌉ | ⌈N5⌉ | ⌈N1⌉ | ⌈N0.1⌉ | max length |
| chr22 reference | 51 324 926 | 16 | 793 | 356 478 | 356 478 | 356 478 |
| chr22 iEFG | 86 646 569 | 23 | 277 | 13 097 | 356 478 | 356 478 |

#### 4.2 Experimental test: comparison to variation graphs

We compare the iEFG obtained in the previous section with the output of the vg toolkit [7]. Using the same T2T-CHM13 reference and 1KGP phased variations, vg takes approximately 10 minutes and 600 MB of space to obtain the graph. On the other hand, if the input is the MSA generated by vcf2multialign, vg takes approximately 1 day and 300 GB of memory[2].

The features of the resulting graphs are compared to those of the iEFG in Table 2. Notably, the vg graphs compress the input more than the iEFG and they have more nodes and edges: the iEFG has 86M total nucleotides compared to the 53M of the vg graphs, i.e. a 63% increase. The width, i.e. the minimum number of paths needed to cover the nodes of the graph, is also significantly higher for the iEFG, which will affect the tools exploiting this parameter such as GraphChainer and minichain. We also study the topological features of the graphs that are independent of how linear unbranching paths are segmented: the iEFG features a reduced amount of recombination due to the constraints imposed by the indexing property (Observation 1). Indeed, the total number of paths in the iEFG that are maximal, that is, they start at a source and end at a sink, is in the order of magnitude of $10^{267\,171}$, compared to that of the vg graph built from the VCF variations, which is equal to $10^{401\,730}$.

Table 2: Features of the chromosome 22 graphs generated from the T2T-CHM13 reference and the 1KGP phased variations. Height H is defined as the largest number of nodes in an iEFG block; width is the size of the smallest set of paths covering the graph nodes; choices is the cumulative degree of branching nodes; branching factor is the largest number of branching nodes encountered in any path; paths is the total number of maximal paths expressed in the graph, represented in scientific notation rounded to the 2nd decimal.

| Method | nodes | edges | bases | N50 | longest node | H | width |
|---|---|---|---|---|---|---|---|
| iEFG | 3 659 566 | 5 054 220 | 86 646 569 | 23 | 356 478 | 543 | 568 |
| vg (VCF) | 4 839 050 | 6 408 323 | 53 073 716 | 32 | 32 | – | 28 |
| vg (MSA) | 4 787 254 | 6 145 861 | 52 782 038 | 32 | 32 | – | 9 |

| Method | branching nodes | choices | branching factor | paths |
|---|---|---|---|---|
| iEFG | 865 808 | 2 260 463 | 841 802 | $4.72 \cdot 10^{267\,171}$ |
| vg (VCF) | 1 423 765 | 2 993 039 | 1 307 293 | $5.19 \cdot 10^{401\,730}$ |
| vg (MSA) | 1 323 783 | 2 682 391 | 1 321 412 | $2.37 \cdot 10^{381\,472}$ |

Then, we evaluate the performance of GraphAligner on the three graphs with the workflow described in the main paper. We obtain three read datasets of 103810, 103395, and 103538 reads from the iEFG, the vg VCF graph, and the vg MSA graph, respectively, and align each dataset to each graph. The accuracy and performance results are shown in Table 3 and Figure 4. There is a clear performance separation in aligning to the iEFG compared to the vg graphs, as using the former generates 5-6 times more seeds and increases the time and space consumed. Even though the sampled vg graph paths spell sequences that are not found in the iEFG, the accuracy of the path and truth ED criteria does not show significant variation except for $\sigma_{\text{truth}} = 0.95$ in the alignment of iEFG reads to the iEFG: we conjecture that nodes from the same iEFG block with high sequence similarity challenge the extending step of GraphAligner.

[2] We disabled the computation of the haplotype paths in the output graph, since vg would otherwise consume all available memory in our machine.

Table 3: Read mapping accuracy of GraphAligner on the chromosome 22 graphs built from the 1KGP-T2T datasets and `BadRead` simulated reads. Cross-aligned reads do not have a ground truth, and thus their corresponding metric information is missing (for $\delta$ and $\sigma_{\text{truth}}$).

| Graph | Reads | Path accuracy | | ED accuracy | |
|---|---|---|---|---|---|
| | | $\delta = 0.1$ | $\delta = 0.95$ | $\sigma_{\text{truth}} = 0.1$ | $\sigma_{\text{read}} = 0.1$ |
| iEFG | iEFG | 95.38 | 83.70 | 94.07 | 86.33 |
| | vg (VCF) | – | – | – | 84.57 |
| | vg (MSA) | – | – | – | 84.80 |
| vg (VCF) | vg (VCF) | 95.95 | 90.63 | 94.90 | 87.07 |
| | iEFG | – | – | – | 87.16 |
| | vg (MSA) | – | – | – | 86.82 |
| vg (MSA) | vg (MSA) | 95.98 | 90.66 | 94.91 | 86.82 |
| | iEFG | – | – | – | 87.13 |
| | vg (VCF) | – | – | – | 86.80 |

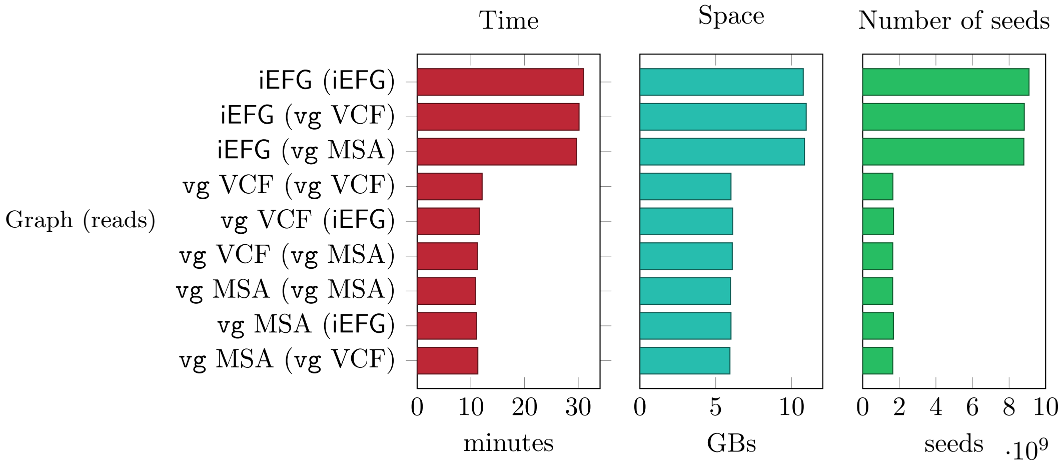

Figure 4: Comparison of the running time, space usage, and total number of seeds of `GraphAligner` using the chromosome 22 iEFG, vg VCF graph, and vg MSA graph. The reads are simulated from the graphs by sampling a random path and simulating long reads with `BadRead`.

### 5 Exact pattern matching, unique seeds

As presented in the main document, iEFGs admit linear-time exact pattern matching by indexing (a variation of) the following string spelled by the concatenation of adjacent nodes [16]:

$$T_{\text{edges}} = \# \cdot \prod_{(u,v)\in E} \ell(uv)\#, \tag{1}$$

where by construction each position $j \in [1..|T_{\text{edges}}|]$ such that $T_{\text{edges}}[j] \neq \#$ corresponds to position $i$ in $\ell(uv)$ for exactly one edge $(u, v) \in E$ and vice versa. In this section, we give a simplified solution that has a slower worst-case running time but is amenable to a practical implementation that extends to a new seeding technique. In particular, we require constant-time support for only the two following operations:

1. LeftExtend$(T_{\text{edges}}, [\ell..r], c)$, given $c \in \Sigma$ and the suffix array interval $[\ell..r]$ of some string $S$ for $T_{\text{edges}} \cdot \$$, returns the suffix array interval of $c \cdot S$ if it occurs in $T_{\text{edges}}$, otherwise the empty interval;

2. edgelocate$(T_{\text{edges}}, \ell)$, given a suffix array index $\ell$ for $T_{\text{edges}} \cdot \$$ corresponding to a suffix of $\ell(uv)$, $(u, v) \in E$, starting at position $i$ in such suffix, returns triple $(u, v, i)$.

The exact pattern matching algorithm is outlined in Algorithm 1 and its main strategy is visualized in Figure 5: a first search finds the longest suffix $Q[f..]$ such that $Q[f..]$ is a prefix of some edge $\ell(uv)$, $(u, v) \in E$; a second search finds an occurrence of $Q[1..f-1]$ in $G$, on the condition that this occurrence of $Q[1..f-1]$ must end at a node boundary; a final search connects $Q[1..f-1]$ to $Q[f..]$ by testing for a matching vertex between the occurrences of $Q[1..f']$ and those of $Q[f'+1..]$, for $f' \in [f..|Q|]$. Since the technical details are complex, we refer the reader to Appendix B for the complete description of the solution and the proof that these searches correctly answer exact pattern matching.

**Algorithm 1:** High-level overview of exact pattern matching of a pattern $Q$ in iEFG $G$ in $O\big(|Q| + \min(|Q|, \mathrm{L}(G))^2 + \mathrm{H}(G)^2\big)$ time after $O(|T_{\text{edges}}|)$-time preprocessing. A full description of the subroutines is given in Appendix B.

**Input:** An EFG $G = (V, E, \ell)$ and query string $Q \in \Sigma^+$
**Output:** Match $(i, P, j)$ of $Q$ in $G$ if it exists, otherwise **false**

Compute $T_{\text{edges}} = \# \cdot \prod_{(u,v)\in E} \ell(uv)\#$;
Preprocess $T_{\text{edges}}$ for backwards pattern matching and edge locate queries;
Execute subroutine `F` for the first search and store result in $f$;
Execute subroutine `S` for the subsequent searches and store result in $P$, $i$, and $y$;
Execute subroutine `C` for the connection and store result in $P'$ and $j$;
**return** $(i, P \cdot P', j)$;

**Theorem 2.** *Let $G = (V, E, \ell)$ be an iEFG. There is an algorithm that, after a preprocessing of $G$ in $O\big(|T_{edges}|\big)$ time (Equation (1)), answers whether $Q \in \Sigma^+$ occurs in $G$ in $O\big(|Q| + \min(|Q|, \mathrm{L}(G))^2 + \mathrm{H}(G)^2\big)$ time, where $\mathrm{L}(G)$ is the longest node length and $\mathrm{H}(G)$ is the maximum block size. In the positive case, the algorithm also reports a match of $Q$ in $G$.*

We implemented exact pattern matching and the greedy seeding algorithm finding semi-repeat-free seeds (srf seeds) described in the main text in our tool `efg-locate`. If multiple threads are

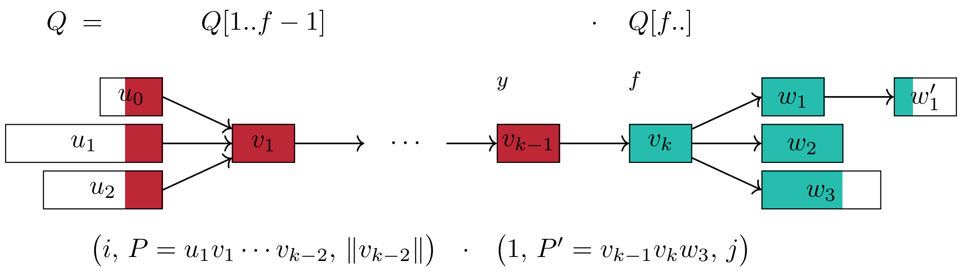

Figure 5: Illustration of the occurrences of a long pattern in an iEFG and of the pattern matching executed by Algorithm 1: the first search finds position $f$, and corresponds to the teal nodes; the second search, corresponding to the yellow nodes, finds an occurrence of $Q[1..f-1]$ (omitting its last node $v_k - 1$) and position $y$; the final step connects the two searches and finds a compatible occurrence of $Q[y..]$.

used, parallelism is offered through the standard method of processing the reads in parallel. We also propose an alternative seeding scheme for finding additional seeds by relaxing the uniqueness constraint of srf seeds as follows. Given a user-defined integer parameter $m \geq 0$, `efg-locate` finds semi-repeat-free seed anchors and, when the search fails in the interval $Q[x..y]$, instead of reporting no match it reports anchors $([x'..y], (i, uv, j))$ with $x'$ the maximum value in $[1..y]$ such that $Q[x'..y]$ occurs at most $m$ times in $T_{\text{edges}}$, if such anchors exist, and we call these anchors srf + edge-$m$.

##### 5.1 Experimental test: srf-based seeds

We compare the use of srf seeds to the default minimizer seeds of `GraphAligner` on the chromosome 22 iEFG from Section 4.1 and the read dataset from Section 4.2, under the experimental setup described in the main paper. The results are illustrated in Figure 6 and Table 4. Out of the 103 810 simulated reads, `efg-locate` finds seeds for the forward or reverse complement strand of 101 697 reads, i.e. 98 % of the total, and the alignment is 7 times faster than `GraphAligner` and reaches a path accuracy of 86.87% for $\delta = 0.1$. Moreover, the number of minimizer seeds found by `GraphAligner` is 250x the number of semi-repeat-free seeds, proving that `efg-locate` finds high-quality seeds in the iEFG.

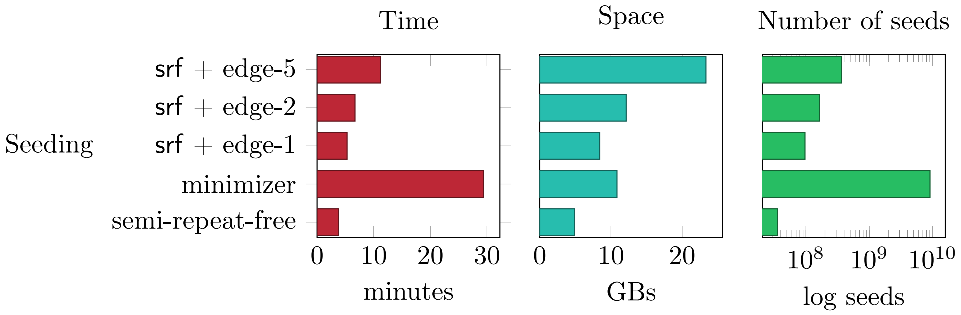

Figure 6: Comparison of running time, space usage, and number of seeds of the read mapping by `GraphAligner` and our aligners based on srf seeds, for the chromosome 22 iEFG constructed from the 1KGP-T2T dataset. Note that for the number of seeds the x-axis does not start at 0.

Table 4: Read mapping accuracy of GraphAligner and our aligners based on srf-based seeds on the chromosome 22 iEFG built from the 1KGP-T2T dataset.

| Seeding technique | Path accuracy | | ED accuracy | |
|---|---|---|---|---|
| | $\delta = 0.1$ | $\delta = 0.95$ | $\sigma_{\text{truth}} = 0.1$ | $\sigma_{\text{read}} = 0.1$ |
| semi-repeat-free | 86.87 | 75.62 | 84.10 | 77.69 |
| GraphAligner | **95.38** | **83.70** | **94.07** | **86.33** |
| srf + edge-1 | 92.36 | 80.79 | 92.11 | 84.10 |
| srf + edge-2 | 92.44 | 80.84 | 92.43 | 84.34 |
| srf + edge-5 | 94.02 | 82.38 | 93.10 | 85.17 |

#### 5.2 Experimental test: maximal exact matches

We compare the use of maximal exact match (MEM) seeds computed by tool `efg-mems` [15] to semi-repeat-free and minimizer seeds on a SARS-CoV-2 iEFG built from the same 100 strains used in the experiments of [15]. The iEFG obtained from `founderblockgraph` presents unique node labels that are too short for the minimizer index of `GraphAligner` under the default settings, thus we merge blocks containing linear paths, a post-processing of the graph which is similar to `vg unchop` but that retains the EFG topology.

As a proof-of-concept, we implemented the mapping of MEMs from the indexes used by `efg-mems` to the nodes of the graph and we wrapped `efg-mems` in a script that considers the reads and their reverse complement for the seed phase, finds MEMs of length at least 20, and stores the intermediate seeds on disk. We also implemented the handling of `N`s in `efg-locate`. Thus we modify the workflow of the main paper by running every aligner using a single compute thread, since `efg-mems` does not support parallel processing of the reads, and we set the read simulation for 1000x coverage, obtaining 2722 reads. The alignments under minimizer, semi-repeat-free, and MEM seeds took 33, 58, and 16433 seconds, respectively; 21, 56, and 1278 kB of space; and obtained 3 762 450, 601770, and and 2 698 999 seeds. The accuracy results are shown in Table 5: there is no big difference in the quality of the seeds. Since semi-repeat-free seeds are 498x faster to find than MEMs, the experiments in the rest of the paper use semi-repeat-free seeds computed by `efg-locate` as the seeding technique.

Table 5: Read mapping accuracy of `GraphAligner` and our aligners based on semi-repeat-free seeds or maximal exact match (MEM) seeds computed by `efg-locate` and `efg-mems -k 20` on a SARS-CoV-2 iEFG built from 100 strains.

| Seeding technique | Path accuracy | | ED accuracy | |
|---|---|---|---|---|
| | $\delta = 0.1$ | $\delta = 0.95$ | $\sigma_{\text{truth}} = 0.1$ | $\sigma_{\text{read}} = 0.1$ |
| minimizers | **70.83** | **51.29** | **81.96** | **66.90** |
| semi-repeat-free | 70.43 | 50.96 | 81.63 | 66.86 |
| MEMs of length $\geq 20$ | **70.83** | **51.29** | **81.96** | **66.90** |

### 6 Chaining on Elastic Founder Graphs

We show how to chain a set of exact match node anchors between a read and an iEFG by adapting the sequence-to-sequence co-linear chaining formulation introduced by Jain et al. [8]. We dedicate this section to explain how the work of Jain et al. [8] works in the sequence-to-sequence case, and our specific modifications to adapt this solution for the sequence-to-iEFG case. First, we define the chaining problem between *two sequences.*

**Definition 2** (Co-linear chain [8]). *Given strings* $Q_p, Q_q \in \Sigma^+$*, let* $A_p = ([x_p..y_p], [a_p..b_p])$*,* $A_q = ([x_q..y_q], [a_q..b_q])$ *be two exact match anchors between* $Q_p$ *and* $Q_q$*. Interval* $[x..y]$ precedes *interval* $[x'..y']$ *if* $x \leq x'$ *and* $y \leq y'$*. We say that* $A_p$ precedes $A_q$ *if both* $[x_p..y_p]$ *precedes* $[x_q..y_q]$ *and* $[a_p..b_p]$ *precedes* $[a_q..b_q]$*. In this case,* $A_p$ *and* $A_q$ *are said to be* co-linear*. A sequence of anchors* $A_1, \dots, A_c$ *is called a* (co-linear) chain *if* $A_p$ *precedes* $A_{p+1}$ *for all* $p \in [1 \dots c-1]$*. The* cost *of a chain* $A_1, \dots, A_c$ *is* $\sum_{p=1}^{c-1} \mathrm{connect}(A_p, A_{p+1})$*, where* connect *is defined as* $\mathrm{connect}(A_p, A_{p+1}) = \mathrm{g}(A_p, A_{p+1}) + \mathrm{o}(A_p, A_{p+1})$ *with*

$$\mathrm{g}(A_p, A_{p+1}) = \max\big(0, x_{p+1} - y_p - 1, a_{p+1} - b_p - 1\big), \qquad \textit{(gap cost)}$$

$$\mathrm{o}(A_p, A_{p+1}) = \big|\max\big(0, y_p - x_{p+1} + 1\big) - \max\big(0, b_p - a_{p+1} + 1\big)\big|. \qquad \textit{(overlap cost).}$$

Two chain-consecutive co-linear anchors $A_p$ and $A_{p+1}$ induce a cost equal to the sum of the maximum gap between them plus the difference of overlaps between them, which is encapsulated into the function $\mathrm{connect}(A_p, A_q)$. The cost of the corresponding chain is, naturally, the total sum of the connect values of anchors consecutive in the chain. In [8], the authors prove that the minimum cost of a chain is equal to the edit distance supported by the anchors[3], thus establishing an elegant connection between edit distance and co-linear chaining. They also show that such a minimum cost chain can be computed by a simple dynamic program in $O(n^2)$ time, where $n$ is the number of anchors: for each anchor $A_q$, compute the minimum cost $C[q]$ of a chain ending with anchor $A_q$, by taking the minimum of $C[p] + \mathrm{connect}(A_p, A_q)$ over all anchors $A_p$ preceeding $A_q$. Moreover, they solve this problem in $O(n \log^4 n)$ time using 4-dimensional data structures and they propose a simpler and practical algorithm that restricts the dynamic programming search to anchors at distance at most $B$ in one of the strings, a parameter which is initially set to the initial guess $B_1$. Then, since the optimal solution might contain consecutive anchors at distance more than $B_1$, at each iteration the algorithm updates $B$ to $B \cdot \alpha$, where $\alpha > 1$ is a given ramp-up parameter. Even though this approach runs in $O(n^2)$ time in the worst case, Jain et al. show that under a uniform and sparse distribution of anchors the algorithm runs in $O(n \cdot B_{\max} + n \log n)$ time, where $B_{\max}$ is the maximum value of $B$ considered. In the conference version of the work by Jain et al. [9], it was initially claimed that the algorithm finds the optimal solution and the solution quality was verified experimentally. Unfortunately, optimality is not always guaranteed [8, Section 5], but the experimental results hold nonetheless. This practical solution was implemented into the tool `ChainX`[4].

Here we adapt `ChainX`'s algorithm for sequence-to-EDS chaining. In our workflow, we apply this exact solution as a heuristic for co-linear chaining on iEFGs, given their similar structure: conceptually, we just add all possible edges between adjacent iEFG blocks to obtain an EDS; we also split each seed spanning multiple nodes into multiple seeds spanning only one node. Even though the EDS chain might not be a valid chain in the iEFG, we can still use the chained anchors to

[3]This specialized version of edit distance is called anchored edit distance in the paper. In particular, the equivalence holds after adding two special initial and final anchors. By changing the connect function for these two special anchors, the minimum cost of a chain can become the semi-global (anchored) edit distance between the string and the graph, which is the objective function used for long reads.

[4]Available at https://github.com/at-cg/ChainX.

obtain high-quality seeds guiding the extension phase. Moreover, this heuristic approach is simpler and faster compared to chaining the original seeds on the iEFG. Additionally, we show that by a small modification of the algorithm by Jain et al. we recover the optimality claim for some of the anchors used in our experiments: instead of checking the distance between the starting positions of anchors, we check for the gap between them. Theorem 3 shows that this modification outputs a minimum chain for anchors that do not overlap on the string.

To adapt `ChainX`, we show how to perform the following two operations in constant time: computing the connect function between anchors, and deciding whether an anchor precedes another in the EDS. Recall the definition of chaining in a graph from the main paper.

**Definition 3** (Co-linear chain on a graph [1]). *Given $Q \in \Sigma^+$ and labeled graph $G = (V, E, \ell)$, let $A_p = ([x_p..y_p], (i_p, u_p, j_p))$ and $A_q = ([x_q..y_q], (i_q, u_q, j_q))$ be exact match anchors between $Q$ and $G$. Then, $A_p$* precedes *$A_q$ if $[x_p..y_p]$ precedes $[x_q..y_q]$ and: $u_p \neq u_q$ and there is a $u_p u_q$-path, or $u_p = u_q$ and $[i_p..j_p]$ precedes $[i_q..j_q]$. The* overlap *in the graph is zero if $u_p \neq u_{p+1}$ and as before otherwise. The* gap *in the graph is defined as before if $u_p = u_{p+1}$ and equal to the shortest sequence length of a $u_p u_{p+1}$-path minus $(j_p + \|u_{p+1}\| - i_{p+1} + 1)$ otherwise. See Figure 7.*

First, to decide if $A_p = \big([x_p..y_p], (i_p, u_p, j_p)\big)$ precedes $A_q = \big([x_q..y_q], (i_q, u_q, j_q)\big)$, we precompute a table $\mathtt{B}$ such that $\mathtt{B}[u] = V_i$ is the block of $u \in V_i$ for each $i \in [1..b]$. For $A_p$ to precede $A_q$, $[x_p..y_p]$ must precede $[x_q..y_q]$. Moreover, if $u_p = u_q$, then $[i_p..j_p]$ must precede $[i_q..j_q]$, and if $u_p \neq u_q$, then $\mathtt{B}[u_p] < \mathtt{B}[u_q]$ must hold. In any other case $A_p$ does not precede $A_q$. For the connect function, we re-use the overlap function of `ChainX` (as overlaps in the EDS can only be computed when $u_p = u_q$) and we compute the gap function as follows. Again, if $u_p = u_q$ then we reuse the gap function from `ChainX`. Otherwise, if $\mathtt{B}[u_p] < \mathtt{B}[u_q]$ we define the gap (in the EDS) as $\|u_p\| - j_p + \sum_{i=\mathtt{B}[u_p]+1}^{\mathtt{B}[u_q]-1} \min_{u \in V_i} \|u\| + (i_q - 1)$, that is the distance from $j_p$ to the end of $\ell(u_p)$, plus the minimum label length of each block inbetween $u_p$ and $u_p$, plus the distance from the beginning of $\ell(u_q)$ until $i_q$ (corresponding graph distance when the EDS is interpreted as a graph). Note that this value can be computed in constant time by precomputing values $\mathtt{S}[1..b]$ such that $\mathtt{S}[i] = \sum_{j=1}^{i} \min_{v \in V_j} \|v\|$. See Figure 7 for an example. Algorithm 2 shows the pseudocode summarizing our adaptation of `ChainX`.

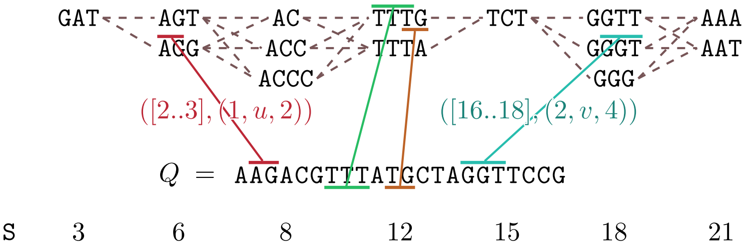

Figure 7: An EDS $G = (V, \ell)$ partitioned into blocks $V_1, \ldots, V_b$, a pattern $Q$, and a chain of co-linear node anchors between $Q$ and $G$. The cost of connecting consecutive anchors in the chain, defined as the maximum gap plus the absolute overlap difference, can be computed in constant time by preprocessing $G$ for the constant-time computation of node block indices and distance between two EDS nodes.

**Theorem 3.** *Algorithm 2 correctly computes the optimal chaining costs $C[1..n]$ when there exists an optimal chain whose anchors do not overlap in $Q$.*

*Proof.* Suppose there is an optimal chain $\mathcal{C}$ with cost $c$ and with the maximum gap between any two consecutive anchors $g \leq c$. Suppose by contradiction, that Algorithm 2 reports a minimum

**Algorithm 2:** Co-linear chaining on EDSes.

**Input:** An EDS $G = (V, \ell)$ whose nodes are partitioned into blocks $V_1, \ldots, V_b$, anchors $\mathcal{A} = A_1, \ldots, A_n$ between a query string $Q \in \Sigma^+$ and $G$, parameters $B_1$ and $\alpha$, and tables $\texttt{B} : V \to [1..b]$ and $\texttt{S}[1..b]$, such that $\texttt{B}[u] = i$ iff $u \in V_i$ and $\texttt{S}[i] = \sum_{j=1}^{i} \min_{\texttt{B}[u]=j} \|u\|$.

**Output:** Cost array $C[1..n]$ such that $C[q]$ is the optimal co-linear chaining cost for any ordered subset of $\mathcal{A}$ ending at $A_q$

Sort anchors $\mathcal{A}$ by increasing starting position in $Q$ into array $A'$;
Initialize all values of $C[1..n]$ to $\infty$ and set $B \leftarrow B_1$;
**repeat**
  $o \leftarrow 1$;
  **for** $q \leftarrow 1$ **to** $n$ **do**
    ▷ Increment $o$ while the gap of $A'[o]$ with $A'[q]$ in $Q$ is at most $B$
    $\big([x_q..y_q], (i_q, u_q, j_q)\big) \leftarrow A'[q]$;
    $\big([x_o..y_o], (i_o, u_o, j_o)\big) \leftarrow A'[o]$;
    **while** $x_q - y_o + 1 > B$ **do**
      $o \leftarrow o + 1$;
      $([x_o..y_o], (i_o, u_o, j_o)) \leftarrow A'[o]$;
    **for** $p \in [o..q-1]$ **do**
      $([x_p..y_p], (i_p, u_p, j_p)) \leftarrow A'[p]$;
      ▷ Check if $A'[p]$ precedes $A'[q]$ in $O(1)$ time
      **if** $[x_p..y_p] < [x_q..y_q] \wedge (\texttt{B}[v_p] < \texttt{B}[v_q] \vee (u_p = u_q \wedge [i_p..j_p] < [i_q..j_q]))$ **then**
        ▷ Compute connect$(A'[p], A'[q])$ in $O(1)$ time
        $\texttt{gap} \leftarrow \max(0, x_q - y_p - 1)$;
        **if** $u_p = u_q$ **then**
          $\texttt{gap} \leftarrow \max(\texttt{gap}, i_q - j_p - 1)$
        **else**
          $\texttt{gap} \leftarrow \max(\texttt{gap}, (\|u_p\| - j_p) + \texttt{S}[\texttt{B}[u_q] - 1] - \texttt{S}[\texttt{B}[u_p]] + (i_q - 1))$
        $\texttt{overlap} \leftarrow \max(0, y_p - x_q + 1)$;
        **if** $u_p = u_q$ **then**
          $\texttt{overlap} \leftarrow |\texttt{overlap} - \max(0, j_p - i_q + 1)|$
        $C[q] \leftarrow \min\left(C[q], C[p] + \texttt{gap} + \texttt{overlap}\right)$;
  $B_{\text{last}} \leftarrow B$;
  $B \leftarrow \alpha \cdot B$;
**until** $C[n] \leq B_{\text{last}}$;
**return** $C[1..n]$;

cost of $c' > c$. By looking at the main loop of the algorithm we know that $c' \leq B_{\text{last}}$, but then $g < B_{\text{last}}$, and thus $\mathcal{C}$ should have been considered in the last iteration. □

Some anchor classes used in our experiments—semi-repeat-free seeds and srf + edge-1 seeds from Section 5.1—do not overlap in $Q$ with other co-linear anchors. As such, every minimum cost chain is made of non-overlapping anchors in $Q$ and Theorem 3 provides correctness of using Algorithm 2 on our datasets. Moreover, for these classes of inputs, we recover the average-case running time guarantee, assuming a uniform anchor distribution and $n \leq |Q|$. Indeed, in the experiments of Section 5.1 the number of semi-repeat-free and srf + edge-1 seeds is always much smaller than the pattern length.

**Theorem 4.** *Let $Q \in \Sigma^+$ be a pattern and $G = (V, \ell)$ an EDS preprocessed for constant-time precedence and connect queries. Given $n$ anchors between $Q$ and $G$, we can find an optimal chain of cost $\mathrm{OPT}$ in $O(n \cdot \mathrm{OPT} + n \log n)$ average-case time, assuming that $n \leq |Q|$, the anchors do not overlap in the query, and the anchor endpoints are uniformly distributed.*

*Proof.* We follow the proof strategy of [8]. The $O(n \log n)$ term comes from sorting the $n$ anchors. Consider an iteration of the main loop of Algorithm 2 (lines 4–24) with a fixed guess $B$ of maximum gap cost. To get the $O(n \cdot OPT)$ term, we define the indicator random variable $X_{p,x}$ to be equal to 1 if $y_p = x$.[5] By the assumption of uniform distribution of anchor endpoints we have that $\mathbb{E}(X_{p,x}) = 1/|Q| \leq 1/n$, also by assumption. Let $X_q$ be the number of anchors inspected when processing $A_q$. Since the anchors do not overlap, $X_q \leq \sum_{p=1}^{n} \sum_{x=x_q-B}^{x_q-1} X_{p,x}$. The expected total number of anchor inspected $X$ is then

$$\mathbb{E}(X) \leq \sum_{q=1}^{n} \sum_{p=1}^{n} \sum_{x=x_q-B}^{x_q-1} \mathbb{E}(X_{p,x}) \leq nB$$

Finally, the total number of anchors inspected during all iterations of Algorithm 2 is

$$O\left(B_1 n \left(1 + \alpha + \alpha^2 + \cdots + \alpha^{\lceil \log_\alpha \mathrm{OPT} \rceil}\right)\right) = O(n \cdot \mathrm{OPT}).$$

□

Finally, we implemented `chainx-block-graph` for the co-linear chaining on the EDS relaxation of an iEFG. The tool supports global and semi-global chaining and accepts the following user-defined parameters: the initial guess $B_1$ for the optimal cost and the ramp-up factor $\alpha$. Additionally, a constant initial guess can be replaced with the guess $\beta \cdot (|Q| - c)$, where $c$ is the coverage of read $Q$ by the input seeds.

#### 6.1 Experimental tests

We evaluate the effect of chaining on the semi-repeat-free-based seeds from the experiments of Section 5.1 by using `chainx-block-graph` to find the optimal chain for both strands of each read. For development purposes, chaining is not integrated into tool `efg-locate` and it is its own part of the workflow, which is executed in parallel, so to respect the given number of available compute threads we split them between `efg-locate` and `chainx-block-graph`.

The accuracy and performance results are shown in Figure 8 and Table 6. On one hand, chaining proves to be a good heuristic to greatly reduce the number of seeds—and thus the amount

[5] In [8] the authors use $x_p = x$ in the definition of the indicator variable. We use $y_p = x$ to consider a correct computation of the gap.

of memory used—while retaining the vast majority of correct alignments. On the other hand, the current implementation does not improve accuracy nor running time in general, showing that `GraphAligner`'s heuristics in the extension phase, considering only the most promising seed clusters, are faster than chaining.

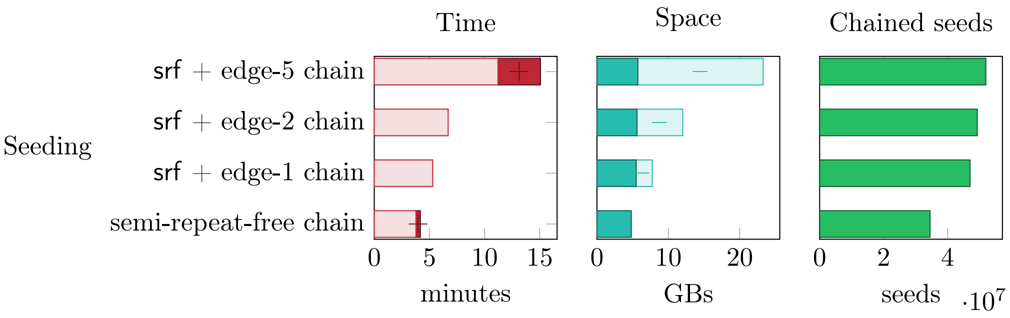

Figure 8: Comparison of running time, space usage, and number of seeds of the read mapping by GraphAligner and our aligners based on *chained* semi-repeat-free seeds, for the chromosome 22 iEFG constructed from the 1KGP-T2T dataset.

Table 6: Read mapping accuracy of GraphAligner and our aligners based on *chained* semi-repeat-free seeds on the chromosome 22 iEFG built from the 1KGP-T2T dataset, compared to the accuracy of Table 4.

| Aligner | Path accuracy | | ED accuracy | |
|---|---|---|---|---|
| | $\delta = 0.1$ | $\delta = 0.95$ | $\sigma_{\text{truth}} = 0.1$ | $\sigma_{\text{read}} = 0.1$ |
| semi-repeat-free chain | (−0.13) 86.74 | (−1.13) 75.49 | (−0.11) 83.99 | (−0.09) 77.60 |
| srf + edge-1 chain | (−1.18) 91.18 | (−1.14) 79.65 | (−1.01) 91.10 | (−0.96) 83.14 |
| srf + edge-2 chain | (−1.32) 91.12 | (−1.28) 79.56 | (−1.17) 91.26 | (−1.12) 83.22 |
| srf + edge-5 chain | (−1.41) 92.61 | (−1.41) 80.96 | (−1.10) 92.00 | (−1.00) 84.17 |

[10] Joseph Lalli. Phased T2T 1KGP panel, February 2023.

[11] Gonzalo Navarro. *Compact data structures: A practical approach.* Cambridge University Press, 2016.

### A Characterization of iEFGs

This section is dedicated to showing that we can check whether a given EFG respects the semi-repeat-free property and is thus indexable: apart from the validation of iEFGs, this is used by the tool `founderblockgraph` to incrementally fix the segmentation when the optional parameter `--heuristic-subset` is used.

**Lemma 2** (Semi-repeat-free characterization)**.** *Let $G = (V, E, \ell)$ be an EFG partitioned into blocks $V_1, \ldots, V_b$. For simplicity, we assume that $G$ is* non-trivial*, in the sense that all nodes present at least one incoming or outgoing edge. Then $G$ is semi-repeat-free if and only if the following property holds: for each vertex $u \in V_i$, $i \in [1..b]$, node label $\ell(u)$ occurs in edge label $\ell(vw)$ for some $(v, w) \in E$ only at position 1, and in such case $v \in V_i$, or at position $\|v\| + 1$, and in such case $w \in V_i$.*

*Proof.* The proof follows by proving the contrapositive statements, that is, that falsifying the antecedent, the semi-repeat-free property, implies that the consequent is false and vice versa. The negation of the property in the statement of this lemma is equivalent to the existence of a vertex $u \in V_i$ such that one of the following holds:

1. $\ell(u)$ occurs in $\ell(vw)$, $(v, w) \in E$, at some position other than 1 or $\|v\| + 1$;
2. it occurs at position 1 in $\ell(vw)$ and $v \notin V_i$ holds; or
3. it occurs at position $\|v\| + 1$ and $w \notin V_i$.

Note that since we assume $G$ to be non-trivial, to satisfy condition 1. it is sufficient to prove that $\ell(u)$ occurs in $\ell(v)$ for some $v \in V$ at some position other than 1, and to satisfy conditions 2. and 3. it is sufficient to prove that $\ell(u)$ is a prefix of $\ell(v)$ with $v \in V_j$ and $j \neq i$.

($\Leftarrow$) If $G$ is not semi-repeat-free, then there exists $u \in V_i$ such that one of the following holds:

- $\ell(u)$ occurs in $G$ starting from the middle of some $v \in V$; then, let $(i, v_1 \cdots v_k, j)$ be a match of $\ell(u)$ in $G$ with $v = v_1 \cdots v_k$ a path in $G$ and $i > 1$; if $k = 1$ then $\ell(u)$ occurs in $\ell(v)$, if $k = 2$ then $\ell(u)$ occurs in $\ell(vv_2)$, and if $k \geq 3$ then $\ell(v_2)$ occurs in $\ell(u)$; all occurrences are at a position different than 1 and condition 1. holds; or
- $\ell(u)$ occurs in $G$ starting from the beginning of some $v \in V_j$ with $j \neq i$; then, let $(1, v_1 \cdots v_k, j)$ be a match of $\ell(u)$ in $G$ with $v = v_1$ and $v_1 \cdots v_k$ a path in $G$; if $k = 1$ then $\ell(u)$ is a prefix of $\ell(v)$ and if $k \geq 2$ then $\ell(v)$ is a prefix of $\ell(u)$; in both cases, conditions 2. and 3. are satisfied.

In both cases the property is falsified, proving that if the property holds then $G$ is semi-repeat-free.

($\Rightarrow$) It is easy to see that any of the conditions 1., 2., and 3. violate the semi-repeat-free property for $G$, since if $\ell(u)$ occurs in $\ell(vw)$ for some $(v, w) \in E$ at position 1 or $\|v\| + 1$ then $\ell(u)$ occurs in $G$ starting from the beginning of node $v \notin V_i$ or $w \notin V_i$, respectively; and that if $\ell(u)$ occurs at some other position in $\ell(vw)$ then $\ell(u)$ occurs in $G$ starting from the middle of $v$ or $w$. ☐

**Theorem 5.** *Given an EFG $G = (V, E, \ell)$ paritioned into blocks $V_1, \ldots, V_b$, we can answer whether it is indexable in $O(|T_{edges}|)$ time (see Equation (1)).*

*Proof.* We can build string $T_{\text{edges}}$ and preprocess it for linear-time pattern matching in $O(|T_{\text{edges}}|)$ time. Additionally, we preprocess $T_{\text{edges}}$ for edge locate queries, as described in Section 5. As already stated in the same section, each position $T_{\text{edges}}[i] \neq \#$ uniquely corresponds to some position $i'$ of $\ell(vw)$, for some $(v, w) \in E$: we can verify the property from Lemma 2 by finding the lexicographical interval $[\ell..r]$ of $\ell(u)$ in the suffix array of $T_{\text{edges}}\$$ for each $u \in V_i$, execute an edge locate query on each $j \in [\ell..r]$, and check whether the property from Lemma 2 is satisfied. If $T_{\text{edges}}$ is built by considering the edges in the order of the blocks $V_1, \ldots V_b$, obtaining the block of any node after the edge locate query can be done in constant time through the standard use of bit vectors indexed for rank and select queries.

However, the complexity of this solution could be greater than $O(|T_{\text{edges}}|)$, since the intervals considered can be nested, so we additionally build the following bit vectors:

- $P[1..|T_{\text{edges}}|+1]$: $P[j] = 1$ if and only if the edge locate query on $j$ corresponds to suffix $\ell(uv)[i'..]$ of $\ell(uv)$ with $i' \neq 1$ and $i' \neq \|u\|+1$;
- $B[1..|T_{\text{edges}}|+1]$: $B[j] = 1$ if and only if the edge locate query on $j$ corresponds to the same block as the result of the edge locate query on $j-1$.

Then, if $[\ell..r]$ is the lexicographic range of $\ell(u)$ in the suffix array of $T_{\text{edges}}$, then the property of Lemma 2 is satisfied if and only if both $B[\ell..r]$ and $P[\ell+1..r]$ contain only 1s. This operation can be answered in $O(1)$ time after the preprocessing of $B$ and $P$ for rank queries, thus proving the statement of this theorem. ☐

### B Exact pattern matching in iEFGs

This section is dedicated to fully describing Algorithm 1 and proving its correctness as stated in Theorem 2, which we repropose here.

**Theorem 2.** *Let $G = (V, E, \ell)$ be an iEFG. There is an algorithm that, after a preprocessing of $G$ in $O\big(|T_{edges}|\big)$ time (Equation (1)), answers whether $Q \in \Sigma^+$ occurs in $G$ in $O\big(|Q|+\min(|Q|, \mathrm{L}(G))^2+\mathrm{H}(G)^2\big)$ time, where $\mathrm{L}(G)$ is the longest node length and $\mathrm{H}(G)$ is the maximum block size. In the positive case, the algorithm also reports a match of $Q$ in $G$.*

Recall that the first search of Algorithm 1, as shown in Figure 5, decomposes $Q$ as $Q[..f-1] \cdot Q[f..]$, where $Q[f..]$ is the largest suffix of $Q$ such that $Q[f..]$ is prefix of $\ell(uv)$ for some edge $(u, v) \in E$. We can indeed prove that all matches of a pattern $Q$ in iEFG $G$ that span many nodes can be decomposed as a match of $Q[1..f-1]$ connected to a match of $Q[f..]$.

**Definition 4** (Parse of a string in a labeled graph)**.** *Let $G = (V, E, \ell)$ be a vertex-labeled graph, with $\ell\colon V \to \Sigma^+$. Given a string $Q \in \Sigma^+$, we define a* parse *of $Q$ in $G$ as any partition $Q_1 \cdots Q_k = Q$ such that there exist node matches $o_1 = (i, u_1, \|u_1\|)$, $o_2 = (1, u_2, \|u_2\|)$, …, $o_k = (1, u_k, j)$ of $Q_1$, …, $Q_k$ in $G$, respectively, such that $o_1 \cdots o_k = (i, u_1 \cdots u_k, j)$ is a match of $Q$ in $G$.*

**Lemma 3** (Semi-repeat-free parsing)**.** *Let $G = (V, E, \ell)$ be an iEFG and let $Q \in \Sigma^+$ be a pattern containing a full node label, i.e. $\ell(v)$ is a substring of $Q$ for some $v \in V$. Then, let $i$ be an occurrence of such $\ell(v)$ in $Q$: for all parses $Q_1 \cdots Q_k$ of $Q$ in $G$ it holds that $Q_{k'} \cdots Q_k = Q[i..]$ for some $k' \in [1..k]$.*

*Proof.* Assume by contradiction that there is some parse $Q_1 \cdots Q_k$ of $Q$ such that $Q_{k'} \cdots Q_k \neq Q[i..]$ for all $k' \in [1..k]$, and let $o_1$, …, $o_k$ be node matches of $Q_1$, …, $Q_k$ in $G$, respectively, as per Definition 4. By hypothesis, there is some $v \in V$ such that $\ell(v)$ occurs in $Q$ at position $i$ and thus $\ell(v)$ is a substring of $Q_1 \cdots Q_k$. Then $\ell(v)$ is a prefix of $Q_{k'} \cdots Q_k$ for some $k' \in [1..k]$, reaching a contradiction, or $\ell(v)$ occurs in $G$ starting inside some node involved in match $o_1 \cdots o_k$, violating the semi-repeat-free property. ☐

**Lemma 4.** *Let $G = (V, E, \ell)$ be an iEFG, let $Q \in \Sigma^+$ be a pattern such that there exists a parse $Q_1 \cdots Q_k$ of $Q$ in $G$ with $k \geq 3$, and let $f \in [1..|Q|]$ be the smallest index of $Q$ such that $Q[f..]$ is* *a prefix of $\ell(uv)$ for some edge $(u,v) \in E$. Then, for all parses $Q_1 \cdots Q_k$ of $Q$ in $G$, it holds that $Q[f..] = Q_{k'} \cdots Q_k$ for some $k' \in [1..k]$.*

*Proof.* Let $Q_1 \cdots Q_k$ be a parse of $Q$ in $G$ with $k \geq 3$ with corresponding node matches $o_1$, $\ldots$, $o_k$ of $Q_1$, $\ldots$, $Q_k$ in $G$, respectively, such that $o = o_1 \cdots o_k$ is a match of $Q$ in $G$, with $o_{k-1} = (i_{k-1}, v_{k-1}, j_{k-1})$ and $o_k = (i_k, v_k, j_k)$. Since $k \geq 3$, by the properties of matches in graphs:

- $Q$ must contain a full node label,
- $o_{k-1} = (1, v_{k-1}, \|v_{k-1}\|)$, and
- $Q_{k-1} = \ell(v_{k-1})$.

As a first consequence of this $f$ is well defined, since $Q_{k-1}Q_k$ is a prefix of $\ell(v_{k-1}v_k)$.

Then, we can show that $Q[f..]$ contains a full node label as a prefix: if this is the case then the thesis of this lemma follows by applying Lemma 3. Assume by contradiction that $\ell(u)$ is not a prefix of $Q[f..]$ for all $u \in V$. By definition of $f$, $Q[f..]$ is a prefix of edge label $\ell(\overline{uv})$ for some $(\overline{u}, \overline{v}) \in E$, but if $|Q[f..]| \geq |\ell(\overline{u})|$ we immediately reach a contradiction, so we assume that $|Q[f..]| < |\ell(\overline{u})|$ and that $Q[f..]$ is a proper prefix of $\ell(\overline{u})$. Consider $Q[f..]$ and $Q_{k-1}Q_k$, since both are suffixes of $Q$, and the relation between their lengths:

- if $|Q[f..]| < |Q_{k-1}Q_k|$ then we contradict the definition of $f$, since $Q_{k-1}Q_k$ is a larger suffix of $Q$ that is prefix of $\ell(v_{k-1}v_k)$, with $(v_{k-1}, v_k) \in E$;
- if $|Q[f..]| = |Q_{k-1}Q_k|$ then $\ell(v_{k-1})$ is a prefix of $Q[f..]$, reaching a contradiction;
- if $|Q[f..]| > |Q_{k-1}Q_k|$ then $\ell(v_{k-1})$ occurs in $G$ starting inside $\overline{u}$, since $Q_{k-1}Q_k$ is a proper suffix of $Q[f..]$, $Q_{k-1} = \ell(v_{k-1})$, and we are already assuming that $Q[f..]$ is a proper prefix of $\ell(\overline{u})$, contradicting the semi-repeat-free property for $v_{k-1}$.

Thus $Q[f..]$ is well-defined and contains a full node label as a prefix, so we can apply Lemma 3 and for all parses $Q_1 \cdots Q_k$ of $Q$, we have that $Q[f..] = Q_{k'} \cdots Q_k$ for some $k' \in [1..k]$. □

**Corollary 1.** *Given iEFG $G = (V, E, \ell)$, query $Q \in \Sigma^+$, and after an $O(|T_{edges}|)$-time preprocessing of $T_{edges}$ for backward pattern matching and edge locate queries, we can answer whether $Q$ has a match $(i, v_1 \cdots v_k, j)$ in $G$ with $k \leq 2$ in $O(|Q|)$ time. In the positive case, we can report such a match, and in the negative case, we can report the smallest index $f \in [1..|Q|]$ such that $Q[f..]$ is a prefix of $\ell(uv)$ for some $(u,v) \in E$ if such index exists.*

*Proof.* Since $T_{\text{edges}}$ is defined in Equation (1) as the concatenation of the edge labels of $G$ separated by a unique character #, the standard backward search of $Q$ in $T_{\text{edges}}$ can locate one occurrence of $Q$ in $\ell(uv)$ for some $(u,v) \in E$, if such occurrence exists. Note that if $G$ has only one block, or if it is built from an MSA with long runs of gaps, there might be nodes $u \in V$ without outgoing and incoming edges that are not represented at all in $T_{\text{edges}}$. For completeness and simplicity, before computing $T_{\text{edges}}$:

- we add a *supersource* $s$ to $V$;
- we connect $s$ to all other *sources* $\hat{v} \in V$ of $G$, i.e. nodes $\hat{v}$ such that no $(u, \hat{v}) \in E$ exists, by adding $(s, \hat{v})$ to $E$;
- we assign label $\mathtt{N}$ to $\ell(s)$, with $\mathtt{N} \notin \Sigma$ a symbol not used in $Q$.

We can combine the search of $Q$ in $T_{\text{edges}}$ with the search for the longest suffix of $Q$ that is prefix of some $\ell(uv)$ by testing if the considered suffixes of $Q$ are preceded by the separator character # in $T_{\text{edges}}$ and report the longest one, as implemented in Algorithm 3. Lemma 4 also guarantees that we can answer the pattern-matching query negatively if we do not fully consume $Q$ and no such index $f$ exists since we rule out that $Q$ has a parse $Q_1Q_2$ in $G$. □

**Algorithm 3:** Subroutine of Algorithm 1 answering whether $Q$ has any match in $G$ spanning at most an edge and computing the longest suffix $Q[f..]$ of $Q$ such that $Q[f..]$ is prefix of some edge label $\ell(uv)$ with $(u, v) \in E$. For simplicity: we assume that operation LeftExtend($T_{\text{edges}}, [\ell..r], Q[0]$) is valid and returns an empty interval; we add a supersource $s$ to $G$ with $\ell(s) = \mathtt{N}$ and connect it to all other sources of $G$.

```
Subroutine F()
    y ← |Q|;
    f ← |Q| + 1;
    [ℓ..r] ← [1..|T_edges| + 1];
    while y ≥ 0 and LeftExtend(T_edges, [ℓ..r], Q[y]) is non-empty do
        [ℓ..r] ← LeftExtend(T_edges, [ℓ..r], Q[y]);
        if LeftExtend(T_edges, [ℓ..r], #) is non-empty then
            f ← y;
        y ← y − 1;
    if y = −1 then                                   ▷ Q occurs in some edge of G
        (u, v, i) ← edgelocate(T_edges, ℓ);
        if i > ‖u‖ then                              ▷ Q occurs in ℓ(v)
            return (i − ‖u‖, v, i − ‖u‖ + |Q| − 1);
        else if i + |Q| − 1 ≤ ‖u‖ then               ▷ Q occurs in ℓ(u)
            return (i, u, i + |Q| − 1);
        else                                         ▷ Q occurs in ℓ(uv)
            return (i, u · v, i + |Q| − 1);
    if f > |Q| then                                  ▷ Q has no match in T_edges nor in G
        return false
    Store f;
```

As shown by Lemma 4, if $Q$ occurs in $G$ and has a parse $Q_1 \cdots Q_k$ with $k \geq 3$, then suffix $Q[f..]$ breaks down all possible matches of $Q$ as a match of $Q[1..f-1]$ followed by a match of $Q[f..]$. Note that $Q[1..f-1]$ is the empty string when $f = 1$, but this condition is already handled in Algorithm 3 so from now on we assume that $f \geq 2$ and $Q[1..f-1]$ is a proper prefix of $Q$. The objective of the second search is then to assess whether $Q[1..f-1]$:

1. has a match in $G$ of the form $(i, u, \|u\|)$, and in such case report nothing, or
2. has a match in $G$ of the form $(i, u_1 \cdots u_k, \|u_k\|)$ with $k \geq 1$, and in such case report the match $(i, u_1 \cdots u_{k-1}, \|u_{k-1}\|)$.

Note that the two cases are mutually exclusive: otherwise, $\ell(u_k)$ from case 2. would be a proper suffix of $u$ from case 1. and the semi-repeat-free property would be violated for $u_k$ (Lemma 1). We leave the connection of $(i, u, \|u\|)$ or $(i, u_1 \cdots u_k, \|u_k\|)$ to some match of $Q[f..]$ for later, by additionally storing index $y \in [1..f]$ such that $y = 1$ in case 1. and $y = f - \|u_k\|$ in case 2.

**Lemma 5.** *Given iEFG $G = (V, E, \ell)$ and query $Q' \in \Sigma^+$, after an $O(|T_{edges}|)$-time preprocessing of $T_{edges}$ for backward pattern matching and edge locate queries, we can answer whether $Q'$ has a match $(i, u_1 \cdots u_k, j)$ in $G$ such that $j = \|u_k\|$ in $O(|Q'|)$ time.*

*Proof.* Standard backward pattern matching of $Q$ in $T_{\text{edges}}$ can be modified for our needs, by iteratively searching substrings of $Q'$ matching full edges $\ell(uv)$ with $(u, v) \in E$. Indeed, the backward search of $Q' \cdot \#$ in $T_{\text{edges}}$ considers the suffixes of $Q'$ that are suffixes of some $\ell(uv)$, with $(u, v) \in E$: by checking if each considered suffix $Q'[x..]\#$ is preceded by $\#$ in $T_{\text{edges}}$, we can identify whether $Q'$ contains a full edge label $\ell(uv)$ with $(u, v) \in E$ as its suffix: $u$ and $v$ are unique due to the semi-repeat-free property (see Lemma 1 and Lemma 3). Then, an edge locate query on the occurrence of $Q'[x..] \cdot \#$ identified retrieves $u$ and $v$. The final solution initally sets variable $q$ to $|Q'|$ and:

1. it performs the backward search of $Q'[1..q] \cdot \#$ in $T_{\text{edges}}$, checking at every iteration whether an additional separator character $\#$ can be read;

2. if such $\#Q'[q'.q]\# = \#\ell(uv)\#$ is identified, it retrieves edge $(u, v) \in E$ via an edge locate query of $Q'[q'.q]\#$, it updates $q$ to value $q - \|v\|$, and it repeats step 1.

Steps 1. and 2. are performed until $Q'$ is fully consumed, and in that case an edge locate query retrieves an occurrence of the last $Q'[1..q]$. If any of the backward searches fails, we answer the query negatively. This solution is almost complete, but it misses the matches $(i, u, \|u\|)$ of $Q'$ such that $u \in V$ is a *source* of $V$, since these strings do not correspond to a suffix of some $\ell(v)\ell(w)\#$ in $T_{\text{edges}}$: to avoid this case, we apply the same supersource technique from Corollary 1.

The procedure, implemented in Algorithm 4, takes $O(|Q'|)$ time since each character of $Q'$ is considered at most twice. More precisely, given $G$, $Q \in \Sigma^+$, and $f \in [2..|Q| + 1]$, Algorithm 4 finds a match $(i, u_1 \cdots u_k, \|u_k\|)$ of $Q[1..f - 1]$ and: if $k > 1$ then it reports $(i, u_1 \cdots u_{k-1}, \|u_{k-1}\|)$ and stores value $f - \|u_k\|$ in variable $y$, to indicate that $Q[y..f - 1]$ should be connected to $Q[f..]$ in the final search[6]; if $k = 1$ then it stores value 1 in variable $y$, to indicate that $Q[y..f - 1] = Q[1..f - 1]$ should be connected to $Q[f..]$. The correctness follows from the semi-repeat-free property: due to Lemma 3, all nodes $u_2, \ldots, u_k$ of reported path $(i, u_1 \cdots u_k, \|u_k\|)$ are common to all matches of $Q$ in $G$. ◻

The third and final search is tasked with finding a match of $Q[y..f - 1] \cdot Q[f..]$ in iEFG $G$ using only $T_{\text{edges}}$ and edge locate queries. This problem is essentially a restricted version of pattern matching on iEFGs and admits linear-time solutions after a polynomial-time indexing of $G$: Equi et al. [5] initially solved it by indexing the concatenation of labels $\ell(uvw)$ of paths of length three in $G$; Rizzo et al. [15] improved the indexing to $O(|T_{\text{edges}}|)$ time by indexing a variation of $T_{\text{edges}}$ with complex machinery (in particular, see [15, Figure 2] to understand why this last search requires a complex solution). First, we can prove that this is equivalent to finding a single vertex that connects the two matches, as shown in Figure 5. Then, we show that we can find such connecting vertex in $O(\min(|Q|, \mathrm{L}(G))^2 + \mathrm{H}(G)^2)$ time using only the stated operations.

**Lemma 6** (Vertex connection). *Let $Q \in \Sigma^+$ have no parse $Q_1, \ldots, Q_{k'}$ in iEFG $G$ such that $k' \leq 2$, let $f$ from Lemma 4 be well-defined and such that $f \geq 2$, and let $(i, u_1 \cdots u_k, \|u_k\|)$ be a match of $Q[1..f - 1]$ in $G$. We define $y$ as 1 if $k = 1$, otherwise $y = f - \|u_k\|$. Then $Q$ occurs in $G$ if and only if there exists a vertex $\overline{v} \in V$ such that:*

1. *$\ell(\overline{v})$ is equal to $Q[f..x]$, with $x = f + \|\overline{v}\| - 1$;*

[6] Here there is no ambiguity and we could store $u_k$, since if $Q$ occurs in $G$ (and has no edge occurrence) then $Q[f..]$ starts with a full node label, but we still have to perform the final connect step, so we unify the two cases in one.

**Algorithm 4:** Subroutine of Algorithm 1 answering whether $Q[1..f-1]$ has a match in iEFG $G$ of the form $(i, u_1 \cdots u_k, \|u_k\|)$. If the answer is positive and if $k > 2$, then it returns match $(i, u_1 \cdots u_{k-1}, \|u_{k-1}\|)$ and $y = f - \|u_k\|$; if $k = 1$, it returns $y = 1$. For simplicity: we assume that operation $\mathrm{LeftExtend}(T_{\mathrm{edges}}, [\ell..r], Q[0])$ is valid and returns an empty interval; we add a supersource $s$ to $G$ with $\ell(s) = \mathtt{N}$ and connect it to all other sources of $G$.

```
Subroutine S()
    Initialize P as an empty list;
    y ← −1;
    q_start ← f − 1;
    q ← q_start;
    [ℓ..r] ← LeftExtend(T_edges, [1..|T_edges| + 1], #);
    while q ≥ 1 do
        [ℓ'..r'] ← LeftExtend(T_edges, [1..|T_edges| + 1], Q[q]);
        if [ℓ'..r'] is empty then        ▷ Q[1..f − 1] has no occurrence (i, u_1 ··· u_k, ||u_k||) in G
            return false
        [ℓ..r] ← [ℓ'..r'];
        q ← q − 1;
        if LeftExtend(T_edges, [ℓ..r], #) is non-empty then        ▷ we read a full edge ℓ(uv)
            (u, v) ← edgelocate(T_edges, ℓ);
            if P is empty then
                y ← f − ||v||;
            P.prepend(u);
            q_start ← q_start − ||v||;
            q ← q_start;
            [ℓ..r] ← LeftExtend(T_edges, [1..|T_edges| + 1], #);
    if P is empty then        ▷ Q[1..f − 1] proper suffix of ℓ(uv)
        y ← 1;
        Store y;
    else if q_start > ||P.head|| then        ▷ Q[1..q_start] suffix of ℓ(uv) of length ≥ ||v||
        (u, v, i) ← edgelocate(T_edges, ℓ);        ▷ arbitrary occurrence of Q[1..q_start]
        P.prepend(u);
        Store P, i, y;
    else        ▷ Q[1..q_start] = ℓ(u)
        i ← 1;
        Store P, i, y;
```

2. *$Q[f..]$ is prefix of $\ell(\overline{v}w)$ for some $(\overline{v}, w) \in E$; and*

3. *$Q[y..x]$ is suffix of $\ell(u\overline{v})$ for some $(u, \overline{v}) \in E$.*

*Proof.* ($\Leftarrow$) If $k = 1$, then $Q$ is clearly a substring of $\ell(u\overline{v}w)$, with $u \cdot \overline{v} \cdot w$ a path in $G$. Otherwise $k \geq 2$, $Q[y..x] = \ell(u_k)\ell(\overline{v})$ occurs in $G$, and all of its matches are from the beginning of $u_k$ only due to Lemma 1. Then $u = u_k$, otherwise the semi-repeat-free property would be violated for $u$ or $u_k$, and $(i, u_1 \cdots u_k \cdot \overline{v} \cdot w, |Q| - x)$ is a match of $Q$ in $G$.

($\Rightarrow$) Let $Q'_1, \ldots, Q'_{k'}$ be a parse of $Q$ in $G$ corresponding to some match $(i', u'_1 \cdots u'_{k'}, j')$ as per Definition 4. Due to our first hypothesis we have that $k' > 2$ and we can apply Lemma 4: $Q'_{k''} \cdots Q'_{k'} = Q[f..]$ for some $k'' \in [1..k']$. More specifically, $k'' \in [2..k' - 1]$: by hypothesis, $Q[1..f - 1]$ is non-empty and thus $k'' \geq 2$; $k'' < k'$, otherwise the definition of $f$ would be violated. Then, we can prove that $u'_{k''} \in V$ is the vertex connection:

1. $Q[f..] = Q'_{k''} \cdots Q'_{k'}$ with $k'' < k'$ and thus $\ell(u'_{k''}) = Q[f..x]$ with $x = f + \|u'_{k''}\| - 1$.

2. By the definition of $f$, $Q[f..]$ is prefix of $\ell(vw)$ for some $(v, w) \in E$, but $Q[f..]$ is also equal to $Q_{k''} \cdots Q_{k'}$ with $k'' < k'$. If $k'' = k' - 1$ then $(u'_{k''}, u'_{k'}) \in E$ and we satisfy condition 2. If $k'' < k' - 1$ then $\ell(u'_{k''}u'_{k''+1})$ is a prefix of $Q[f..]$ and by Lemma 1 all occurrences of $Q[f..]$ start from the beginning of $u'_{k''}$, thus $v = u'_{k''}$ and we satisfy the condition as well.

3. If $k = 1$ then $y = 1$ and by hypothesis $(i, u_1, \|u_1\|)$ is a match of $Q[1..f - 1] = Q'_1 \cdots Q'_{k''-1}$ in $G$. In this case, it is easy to see that $k'' = 2$: indeed, if $k'' > 2$ then $\ell(u'_{k''-1})$ is a proper suffix of $\ell(u_1)$, violating the semi-repeat-free property for $u'_{k''-1}$. Otherwise $k > 1$, implying that $y = f - \|u_k\|$. We can prove that $u_k = u'_{k''-1}$: string $Q[y..x] = \ell(u_k)\ell(u'_{k''})$ is a suffix of $Q'_1 \cdots Q'_{k''}$ and occurs in $G$ only from the beginning of $u_k$ due to Lemma 1, so the semi-repeat-free property forces $u'_{k''}$ to be the same node as $u_k$. □

**Corollary 2.** *Given iEFG $G = (V, E, \ell)$ partitioned in blocks $V_1, \ldots, V_b$ and query $Q \in \Sigma^+$, let $Q$ have no match $(i, u_1 \cdots u_k, j)$ in $G$ such that $k \leq 2$. Also, let $f$ from Lemma 4 be well-defined and given in input as well, and let $(i, u_1 \cdots u_k, \|u_k\|)$ be a given match of $Q[1..f - 1]$ in $G$. Then, after an $O(|T_{edges}|)$-time preprocessing of $T_{edges}$ for backwards pattern matching and edge locate queries, we can answer whether $Q$ has a match in $G$ in $O(\min(|Q|, \mathrm{L}(G))^2 + \mathrm{H}(G)^2)$ time, where $\mathrm{L}(G) = \max_{v \in V} |\ell(v)|$ is the longest node length and $\mathrm{H}(G) = \max_{i \in [1..b]} |V_i|$ is the maximum block height of $G$.*

*Proof.* Lemma 6 states that $Q$ occurs in $G$ if and only if there is a vertex $\overline{v} \in V$ making the connection of $Q[1..f - 1]$ with $Q[f..]$ possible. For each $x \in [f..]$, by just using $T_{\mathrm{edges}}$ we can check if $Q[y..x]$ (with $y$ as defined in Lemma 6) contains a full node label $\ell(v)$ as suffix in $O(x - y + 1) = O(|Q|)$ time by querying whether $Q[y..x] \cdot \#$ occurs in $T_{\mathrm{edges}}$ and executing one edge locate query on the relative suffix array interval: the semi-repeat-free property guarantees that if $Q[f..x] = \ell(\overline{v})$ for some $\overline{v} \in V$ then all occurrences of $\ell(\overline{v})$ as suffix of $\ell(uv)$ for some $(u, v) \in E$ are such that $v = \overline{v}$. Thus, if $(u, v, i)$ is the result of the edge locate query on $Q[y..x] \cdot \#$, $Q[f..x] = \ell(v')$ for some $v' \in V$ if $x - y + 1 \geq \|v\|$ (or equivalently $i \leq \|u\|$) and such node $v'$ is exactly $v$. For each $x \in [f..]$ fulfilling this condition and corresponding to $v \in V$, we can test if any occurrence of $Q[f..]$ corresponds to an edge $(v'', w) \in E$ such that $v = v''$. This solution is implemented in Algorithm 5.

The correctness follows from Lemma 6 and the arguments above, and we obtain the stated time complexity as follows:

- for finding good $x$ values, testing all strings $Q[y..x]\#$ takes $O(|Q| \cdot \min(|Q|, L))$ time, but we can ignore values of $x$ such that $y - x + 1 > 2L$ because $Q[y..x]$ cannot be longer than any edge label $\ell(uv)$;

- we can consider edges $(u', v') \in E$ corresponding to $Q[f..]$ only once and only if some full node label is recognized in the previous step, after collecting all good $x$ values in a list of at most $|Q|$ elements.

The former step takes $O(\min(|Q|, \mathrm{L}(G))^2)$ time and the latter $O(\mathrm{H}(G)^2)$. □

---

**Algorithm 5:** Subroutine of Algorithm 1 connecting a given match $(i, u_1 \cdots u_k, \|u_k\|)$ of $Q[1..f-1]$ in iEFG $G$ to some match of $Q[f..]$ in $G$ by finding a connecting vertex as per Lemma 5. If $k = 1$ and thus $y = 1$, the subroutine reports a match of $Q$ in $G$, otherwise $y = f - \|u_k\|$ and it stores match $(1, P', j)$ of $Q[y..]$ in $G$ in variables $P'$ and $j$.

---

**Subroutine** C()
    Initialize list $L_1$ of elements with values in $[f..|Q|]$;
    **for** $x \in [f.. \min(|Q|, y + 2\,\mathrm{L}(G) - 1)]$ **do**
        **if** $Q[y..x]\#$ *occurs in* $T_{\text{edges}}$ **then**
            $L_1 \leftarrow L_1 \cup x$;
    **if** $L_1$ *is empty* **then**
        **return** ***false***
    **for** $\ell \in \mathrm{LeftExtend}(T_{\text{edges}}, [1..|T_{\text{edges}}| + 1], Q[f..])$ **do**
        $(v, w) \leftarrow \mathrm{edgelocate}(T_{\text{edges}}, \ell)$;
        **if** $f + \|v\| \in L_1$ **then** ▷ $v$ is connecting vertex with $\ell(v) = Q[f..x]$
            $x \leftarrow f + \|v\| - 1$;
            $[\ell'..r'] \leftarrow \mathrm{LeftExtend}(T_{\text{edges}}, [1..|T_{\text{edges}}| + 1], Q[y..x])$;
            $(u, v, i) \leftarrow \mathrm{edgelocate}(T_{\text{edges}}, \ell')$; ▷ arbitrary occurrence of $Q[y..x]$
            **if** $y = 1$ **then**
                **return** $(i, uvw, |Q| - f + 1 - \|v\|)$
            **else**
                $P' \leftarrow uvw$;
                $j \leftarrow |Q| - f + 1 - \|v\|$;
                **Store** $P', j$;
    **return** ***false*** ▷ no connecting vertex exists

---

Finally, Theorem 2 is proved by Corollary 1, Lemma 5, and Corollary 2. Algorithms 3 to 5 fully describe Algorithm 1.

### C Full results of experimental tests

Table 7: Uniqueness statistics on the semi-repeat-free segmentations of the T2T-CHM13 human chromosomes.

| chromosome | bases | ⌈N50⌉ | ⌈N5⌉ | ⌈N1⌉ | ⌈N0.1⌉ | max length |
|---|---|---|---|---|---|---|
| chr1 | 248 387 328 | 16 | 153 | 6 212 | 14 839 | 14 839 |
| chr2 | 242 696 752 | 16 | 38 | 575 | 6 523 | 17 928 |
| chr3 | 201 105 948 | 16 | 39 | 1 633 | 8 616 | 14 831 |
| chr4 | 193 574 945 | 16 | 41 | 3 086 | 12 882 | 14 238 |
| chr5 | 182 045 439 | 16 | 41 | 2 127 | 10 984 | 17 752 |
| chr6 | 172 126 628 | 16 | 39 | 3 499 | 17 992 | 19 231 |
| chr7 | 160 567 428 | 16 | 45 | 2 487 | 10 855 | 10 855 |
| chr8 | 146 259 331 | 16 | 39 | 2 203 | 8 165 | 9 483 |
| chr9 | 150 617 247 | 16 | 3 802 | 46 435 | 48 873 | 48 873 |
| chr10 | 134 758 134 | 15 | 41 | 2 027 | 9 624 | 12 184 |
| chr11 | 135 127 769 | 15 | 41 | 2 235 | 15 312 | 17 892 |
| chr12 | 133 324 548 | 15 | 39 | 1 257 | 11 386 | 13 213 |
| chr13 | 113 566 686 | 16 | 363 | 1 579 331 | 1 579 331 | 1 579 331 |
| chr14 | 101 161 492 | 15 | 54 | 7 448 | 298 974 | 298 974 |
| chr15 | 99 753 195 | 16 | 1 330 | 534 092 | 534 092 | 534 092 |
| chr16 | 96 330 374 | 16 | 1 755 | 12 266 | 21 081 | 21 341 |
| chr17 | 84 276 897 | 15 | 70 | 4 593 | 9 522 | 10 360 |
| chr18 | 80 542 538 | 15 | 53 | 6 105 | 16 148 | 16 148 |
| chr19 | 61 707 364 | 16 | 118 | 4 609 | 10 430 | 10 430 |
| chr20 | 66 210 255 | 15 | 52 | 4 016 | 6 806 | 9 898 |
| chr21 | 45 090 682 | 15 | 222 | 825 895 | 825 895 | 825 895 |
| chr22 | 51 324 926 | 16 | 793 | 356 478 | 356 478 | 356 478 |
| chrX | 154 259 566 | 16 | 46 | 1 912 | 10 756 | 12 149 |
| chrY | 62 460 029 | 937 | 8 002 | 34 050 | 40 632 | 40 632 |

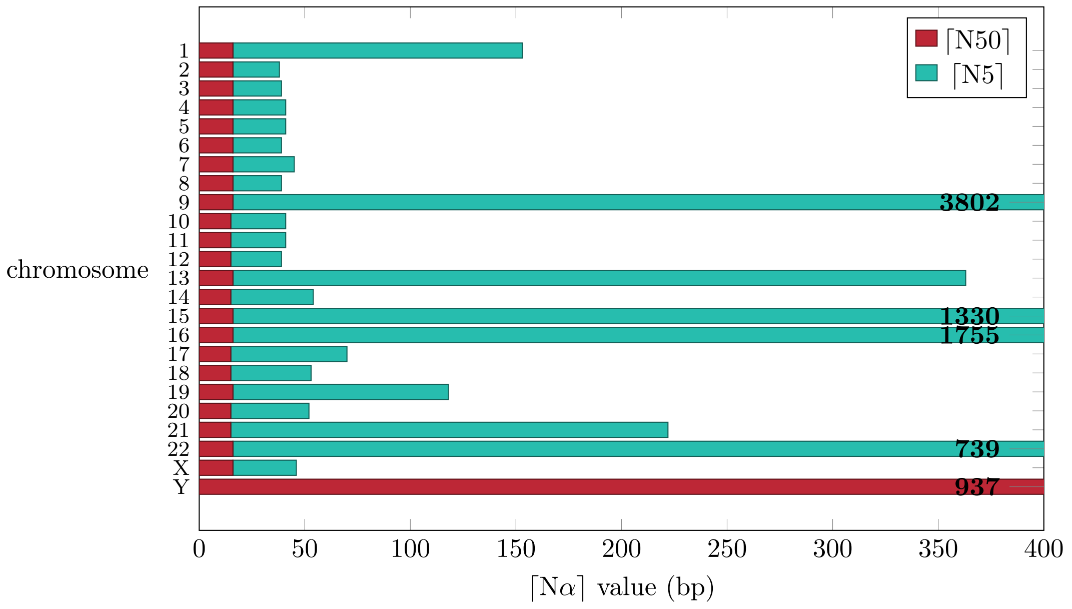

Figure 9: Visualization of the ⌈N50⌉ and ⌈N5⌉ values for the semi-repeat-free segmentations of T2T-CHM13 individual chromosomes.

#### D Commands used in the experiments

- to obtain the chromosome 22 MSA with `vcf2multialign`:

```
vcf2multialign --input-reference=chr22_uppercase.fasta
    --input-variants=1KGP.CHM13v2.0.chr22.phased.vcf.gz
    --chromosome chr22 --haplotypes
    -s msa.fasta
```

- iEFG construction for chromosome 22:

```
founderblockgraph --elastic --gfa --ignore-chars="N" --output-paths
    --threads=64 --heuristic-subset 250
    --input=msa.fasta --output=chr22-efg.gfa
```

- iEFG construction for SARS-CoV-2:

```
founderblockgraph --elastic --gfa --ignore-chars="N" --output-paths
    --threads=64
    --input=msa.fasta --output=SARS-CoV-2-efg.gfa
efg-simplify --simplify-tunnels
    SARS-CoV-2-efg.gfa SARS-CoV-2-efg-simplified.gfa
```

- `vg` construction of the chromosome 22 graph from VCF input:

```
vg construct -t 64 -r chr22_uppercase.fasta
    -v 1KGP.CHM13v2.0.chr22.phased.vcf.gz > vggraph.vg
vg convert -t 64 -f vggraph.vg > vggraph.gfa
```

- `vg` construction of the chromosome 22 graph from MSA input:

```
vg construct -M msa.fasta -p > output/vgmsagraph.vg
vg convert -t 64 -f vgmsagraph.vg > vgmsagraph.gfa
```

- semi-repeat-free ($m = 0$) and srf + edge-$m$ seed computation for `GraphAligner`:

```
efg-locate --approximate --reverse-complement
    --split-output-matches-graphaligner --ignore-chars="N"
    --approximate-edge-match-min-count=m --threads=64
    chr22-efg.gfa reads.fasta seeds.gaf
```

- chaining of seeds in the iEFG (`efg-locate` should be called with option `--split-output-matches` and with an appropriate number of threads for load balancing):

```
chainx-block-graph --semi-global --threads 64
    --split-output-matches-graphaligner
    chr22-efg.gfa seeds.gaf chains.gaf
```

- extending seeds or chains into alignments with `GraphAligner`:

```
graphaligner --max-cluster-extend 5 -b 10 -t 64
     -g chr22-efg.gfa -f reads.fastq
     --realign seeds.gaf -a alignments.gaf
```

- alignment with `minigraph`:

```
minigraph -t 64 -cx lr chr22-efg.gfa reads.fastq -o alignments.gaf
```

- alignment with `minichain`:

```
minichain -t 64 -c chr22-efg.gfa reads.fastq -o alignments.gaf
```

- alignment with `GraphAligner`:

```
GraphAligner -t 64 -g chr22-efg.gfa -f reads.fastq
    -a alignments.gaf
```

- alignment with `GraphChainer`:

```
GraphChainer -t 64 -g chr22-efg.gfa -f reads.fastq
    -a alignments.gaf
```
